# Cytoplasmic physical state governs the influence of oxygen on *Pinus densiflora* seed ageing

**DOI:** 10.1101/2020.12.11.421446

**Authors:** Davide Gerna, Daniel Ballesteros, Wolfgang Stöggl, Erwann Arc, Charlotte E. Seal, Chae Sun Na, Ilse Kranner, Thomas Roach

## Abstract

During desiccation, the cytoplasm of orthodox seeds solidifies in a glass with highly restricted diffusion and molecular mobility, which extend longevity. Temperature and moisture determine seed cellular physical state, and oxygen can promote deteriorative reactions of seed ageing. However, whether seed physical state affects O_2_-mediated biochemical reactions during ageing remains unknown. Here, we answered this question using oil-rich *Pinus densiflora* seeds aged by controlled deterioration (CD) at 45 °C and distinct relative humidities (RHs), resulting in a glassy (9 and 33% RH) or fluid (64 and 85% RH) cytoplasm. Regardless of CD regimes, the cellular lipid domain remained always fluid. Hypoxia (0.4% O_2_) prevented seed deterioration only in the glassy state, limiting non-enzymatic lipid peroxidation, consumption of antioxidants (glutathione, tocopherols) and unsaturated fatty acids, accompanied by decreased lipid melt enthalpy and lower concentrations of aldehydes and reactive electrophile species (RES). In contrast, a fluid cytoplasm promoted faster seed deterioration and enabled the resumption of enzymatic activities implicated in glutathione metabolism and RES detoxification, regardless of O_2_ availability. Furthermore, seeds stored under dry/cold seed bank conditions showed biochemical profiles similar to those of CD-aged seeds with glassy cytoplasm under normoxia. These findings are discussed in the context of germplasm management.

**Highlight:** lipid peroxidation occurred during seed ageing in the glassy state and, like viability loss, could be prevented by hypoxia. Seeds with fluid cytoplasm aged faster and irrespective of oxygen availability.

## Introduction

The preservation of seed viability and quality during storage is at the basis of plant propagation and of primary interest for seed banks in agriculture, forestry, and biodiversity conservation (Colville and Pritchard, 2019; Li and Pritchard, 2009; Whitehouse *et al*., 2020). The extended longevity of desiccation tolerant (i.e. orthodox) seeds under dry and cold conditions critically depends on their ability to tolerate both desiccation to water contents (WCs) lower than 0.1-0.07 g H_2_O g^-1^ dry weight (DW) and sub-zero temperatures (Walters, 2015). At the low WC and temperature of conventional storage in seed banks, the cytoplasm of seeds is stabilised by formation of an intracellular glass (referred to as “glassy state”), resulting from the non-crystalline solidification of the cytoplasmic matrix and the entrapment of all cellular organelles within (Ballesteros *et al*., 2020). The glassy cytoplasm restricts molecular diffusion, decelerating the rates of biochemical reactions implicated in seed deterioration, thus extending longevity (Sun, 1997; Murthy *et al*., 2003; Buitink and Leprince, 2008; Ballesteros and Walters, 2011; Fernández-Marín *et al*., 2013; Walters *et al*., 2005a).

In addition to the well-studied influence of WC and storage temperature [e.g. viability equations; (Ellis and Roberts, 1980)], seed longevity is also affected by the gaseous environment during storage. Early reports describe the advantage of hermetical storage to seed longevity (Harrison and McLeish, 1954; Roberts, 1961), and more recent studies show that elevated O_2_ partial pressure shortens seed longevity (Groot *et al*., 2012; Groot *et al*., 2015; Hourston *et al*., 2020). There is consensus that oxidative reactions, which cause the accumulation of macromolecular damage, occupy a primary position in seed ageing and death (McDonald, 1999; Bailly, 2004; Kranner *et al*., 2006; Rajjou and Debeaujon, 2008; Kranner *et al*., 2010; Walters *et al*., 2010; Kumar *et al*., 2015; Bailly, 2019). In the glassy state, limited molecular motion (Ballesteros and Walters, 2011, 2019) is still compatible with the production of reactive oxygen species (ROS) and the consumption of antioxidants, which influence seed redox state (Oracz *et al*., 2009; Bahin *et al*., 2011; Bazin *et al*., 2011; Nagel *et al*., 2015;).

Under the restricted molecular mobility and diffusion within the glass, ROS-processing enzymes cannot access their substrates in the aqueous domain. Hence, low-molecular-weight (LMW) antioxidants offer the only protection from oxidative damage and include tocochromanols in the seed cytoplasmic lipid domain (e.g. membranes and oil bodies), and glutathione (γ-L-glutamyl-L-cysteinyl-glycine, GSH) and ascorbate (L-threo-hexenon-1,4-lacton or vitamin C, AsA) in the cytoplasmic aqueous domain (Kranner *et al*., 2010). Tocochromanols are amphipathic compounds of the vitamin E family (i.e. tocopherols, tocotrienols, and tocomonoenols), which scavenge peroxyl (i.e. lipid) radicals and thus block the propagation phase of lipid peroxidation (Munné-Bosch and Alegre, 2002; Menè-Saffranè *et al*., 2010). Typically, α- and γ-tocopherols are abundant in seeds, particularly in those rich in oil storage reserves (Smirnoff, 2010; Fernández-Marín *et al*., 2017). Dry seeds mainly contain the tripeptide and LMW thiol GSH, and only traces, if any, of AsA (Colville and Kranner, 2010; Gerna *et al*., 2017; Gerna *et al*., 2018). Both GSH and AsA donate an electron to ROS radicals, subsequently converting to glutathione disulphide (GSSG) and dehydroascorbic acid, respectively (Tommasi *et al*., 2001; Kranner *et al*., 2006). In addition, these two water-soluble antioxidants may also help protect the lipid phase by regenerating tocochromanols from tocopheryl radicals, formed by the scavenging of peroxyl radicals produced during lipid peroxidation (Smirnoff and Wheeler, 2000; Munné-Bosch and Alegre, 2002; Colville and Kranner, 2010). A broad range of bioactive molecules is released from lipid peroxides, depending on the type of fatty acid (FA) and how the peroxide decays. The presence of a carbonyl group confers electrophilicity, which is enhanced when the carbonyl is conjugated to an alkene (forming an α,β-unsaturated carbonyl), as found in the so-called reactive electrophile species (RES) (Farmer and Davoine, 2007; Mano *et al*., 2019). Due to its nucleophilic nature, GSH conjugates with RES through reactions catalysed by various glutathione-S-transferases (GSTs, EC 2.5.1.18) enabling detoxification (Roach *et al*., 2018b; Mano *et al*., 2019). Less reactive aldehydes are converted to carboxylic acids by aldehyde dehydrogenases, using NAD(P)^+^ as a cofactor (Mano, 2012). Importantly, GSH is a major cellular redox buffer in dry seeds, and changes in GSH and GSSG concentrations shift the glutathione half-cell reduction potential (E_GSSG/2GSH_, i.e. the glutathione redox state) towards more negative (i.e. more oxidising) values (Schafer and Buettner, 2001; Kranner *et al*., 2006). An oxidative shift in E_GSSG/2GSH_ has been correlated with seed viability, regardless of ageing regimes (Kranner *et al*., 2006; Birtić *et al*., 2011; Chen *et al*., 2013; Nagel *et al*., 2015; Roach *et al*., 2018a). Nonetheless, the combined effects of changes in molecular mobility and O_2_ availability on GSH metabolism during seed storage, and the potential repercussion on to biochemical changes in the lipid domain, are not clear.

Most studies on the biochemical reactions implicated in seed ageing have been conducted using protocols of controlled deterioration (CD), consisting in seed exposure to high temperature (e.g. 35-45 °C) and elevated relative humidity (RH, e.g. 60-70%), ensuring fast declines of viability (Powell and Matthews, 1981; Hay *et al*., 2008). However, accelerating seed ageing using humid/warm conditions typical of CD does not always lead to the same biochemical changes that occur in dry/cold storage conditions of seed banks (Nagel *et al*., 2015; Roach *et al*., 2018a; Nagel *et al*., 2019). For example, viewed via a lack of changes in FA composition and tocochromanol concentrations, the lipid phase remains relatively stable during CD, even in some cases until viability loss under elevated O_2_ concentrations (Lehner *et al*., 2008; Morscher *et al*., 2015; Roach *et al*., 2018a; Schausberger *et al*., 2019), whereas tocochromanol consumption may occur during cold storage of oily and non-oily seeds (Seal *et al*., 2010a; Seal *et al*., 2010b; Roach *et al*., 2018a). The physical properties affecting molecular mobility under these fast (i.e. CD) and slow (i.e. seed bank) ageing regimes can account for different biochemical responses. During dry and cold storage, the conditions fall below the glass transition temperature (Tg), and the seed cytoplasm is in a solid/glassy state (henceforth referred to as glassy). In contrast, elevated RH combined with high temperatures are typically used during CD and lead to fluidisation of the cytoplasm, which enters a liquid/rubbery state (hereafter referred to as fluid) (Walters, 1998; Walters *et al*., 2010; Ballesteros and Walters, 2011).

In this paper, we provide a deeper insight into the role of O_2_ in seed ageing in both the glassy and fluid state. We tested the hypothesis that O_2_ is detrimental to seed longevity, via promoting lipid peroxidation, only when seeds are in a glassy state with restricted enzyme activity and limited protection against oxidative damage. We chose *Pinus densiflora* (Japanese red pine), a widespread species with oily seed storage reserves, inhabiting coniferous forests in central Asia and of interest for reforestation (Washitani and Saeki, 1986; Hu *et al*., 2020). We treated seeds with CD under normoxia (nominal 21% O_2_) and hypoxia (nominal < 1% O_2_) at various RHs to achieve contrasting intracellular physical properties. These were determined by dynamic mechanical analysis (DMA) and differential scanning calorimetry (DSC), which revealed transitions in the visco-elastics and melting properties of both the aqueous and lipid domains of the cytoplasm (Walters *et al*., 2010; Ballesteros and Walters, 2011, 2019; Porteous *et al*., 2019). Sorption isotherms were constructed and assessed to calculate values of the Brunauer-Emmet-Teller (BET) monolayer, which describes the chemical affinity of a material for water and is expressed as the WC at which all water-binding sites at the adsorbent surface are filled with water molecules. The removal of water from the BET monolayer has been proposed to promote deterioration by exposing macromolecules to O_2_ (Labuza, 1980; Buitink *et al*., 1998; Ballesteros and Walters, 2007b; Barden and Decker, 2016), and here we studied if removing the BET monolayer affected biochemical changes accompanying seed deterioration. To characterise the influence of O_2_ on seed redox biochemistry during ageing, we assessed GSH, GSSG, and tocochromanol concentrations using high-performance liquid chromatography (HPLC), FA profiles with gaschromatography coupled to mass-spectrometry (GC-MS), RES and other aldehydes with ultra HPLC-MS/MS. Furthermore, to clarify whether O_2_-dependent CD-induced processes are representative of long-term cold storage in the glassy state, seeds stored for 20 years under seed bank conditions were also analysed.

## Material and methods

### Seed material and storage conditions

All experiments were conducted using *Pinus densiflora* Sieb. et Zucc. (also known as Japanese red pine) seeds obtained from the National Baekdudaegan Arboretum (Seobyeok-ri, Chungyang-myeon, Bonghwa-gun, South Korea). In autumn 2015, seeds were harvested from individual trees in the Gwangneung forest and randomly pooled together. Thereafter, seeds were equilibrated at 30 ± 1.5% RH for about seven weeks and kept until 2019 at −20 ± 2 °C and 56 ± 7% RH, measured with dataloggers (EasyLog, Lascar Electronics Ltd, Whiteparish, UK), in vacuum-sealed laminated polyamide/polyethylene bags. These seeds were used as “control” and had a WC of 0.04 g H_2_O g^-1^ DW before equilibrating to the WCs used during the various CD regimes. In addition, a historic collection of seeds, harvested in 1999 from the same location as 2015 with a total germination of 91% in 1999 and kept inside laminated plastic bags at 4 °C and 0.06 g H_2_O g^-1^ DW for 15 years (hereafter referred to as “seed bank” seeds), was available and was included in the study. In 2015, these seed bank seeds were transferred to −20 °C until analyses in 2019.

### CD and germination

Approximately 4.5 g of seeds were collected in Manila hemp-cellulose bags (Jeden Tag, Zentrale Handelgesellschaft GmbH, Offenburg, Germany) and sealed in 1-L glass jars containing 50 mL of LiCl solutions at 8.6 ± 0.4, 32.9 ± 1.0, 63.9 ± 1.6, and 84.9 ± 1.7% RH and data loggers (EasyLog, Lascar Electronics Ltd, Whiteparish, UK) to monitor temperature and RH during storage. For each replicate, the bags containing seeds were placed in separate jars and incubated at room temperature (RT) in the dark for pre-equilibration to the various RH (Supplementary Table S1 at *JXB* online). During the preequilibration period, sample fresh weights (FWs) were recorded daily and, once they had stabilised over two consecutive days, the jars were flushed with N_2_ to establish hypoxia. This was defined *a priori* as O_2_ concentrations < 1% inside the jars, detected with oxygen sensor spots (PSt3) inside the glass jars in conjunction with a fibre optic O_2_ meter (Fibox 3, PreSens Precision Sensing GmbH, Regensburg, Germany). Subsequently, seeds were further equilibrated at RT in the dark for two days, before starting CD at 44.5 ± 0.4 °C under normoxia (19.6 ± 1.5% O_2_) and hypoxia (0.4 ± 0.5% O_2_). At regular intervals during CD, O_2_ concentrations of all replicates were monitored, while keeping jars at 44.5 ± 0.4 °C. Details on the CD regimes, including duration of individual treatments, average temperature, RH, and O_2_ concentration, and seed WC values are summarised in Supplementary Table S1.

The design of CD experiments aimed at elucidating the effects of O_2_ depletion on viability, biophysical, and biochemical changes between seeds aged for the same duration at the same temperature and RH, targeting a 50% viability loss (P50) under normoxia only. This approach allowed comparisons between seeds subjected to CD with the same physical state but under normoxia or hypoxia. Pilot CD experiments were conducted at about 30, 60, 80, and 100% RH and 45 °C under normoxia to define the duration of CD intervals to reach P50. At least three intervals for each RH were used to estimate the P50 values of seeds aged at all pre-tested RHs via probit analysis (Ellis and Roberts, 1980). At 9% RH, P50 was predicted by plotting the experimental P50 values at 30, 60, 80, and 100% RH against their corresponding WCs (Supplementary Fig. S2). Based on the CD pilot studies, seed viability was assessed by scoring total germination after different CD intervals, depending on RH (Supplementary Table S1). Fifty seeds per replicate were sown in Petri dishes containing three layers of filter paper (Whatman grade 1, GE healthcare, Little Chalfont, United Kingdom) and imbibed with 4 mL of ultrapure water (UPW), prior to germination at 20 °C with a 14 h day (47 ± 3 μmol m^-2^ s^-1^): 10 h night photoperiod. A seed was considered germinated when radicle length exceeded seed length. Scoring total germination ceased when microbial contamination led to first signs of seed decomposition, generally two weeks after the last seed had germinated. The effects of CD under normoxia and hypoxia on germination speed, a proxy for seed vigour, were estimated by calculating the time to reach 25% germination (T_25_) according to the following equation adapted from (Farooq *et al*., 2005):

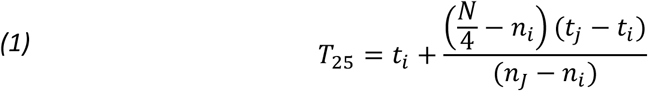

where *N* is the total number of seeds per replicate, n_j_ and n_i_ the cumulative numbers of seeds germinated between consecutive scorings at time t_j_ and t_i_, when n_i_ < N/4 < n_j_.

### Biophysical analyses

#### Dynamic Mechanical Analysis

DMA was conducted to measure structural relaxations and determine the Tg of *P. densiflora* seeds based on the visco-elastic properties of their cytoplasm (Ballesteros and Walters, 2011, 2019). DMA and not DSC was selected due to higher sensitivity to detect the Tg of dry seeds (Ballesteros and Walters, 2011, 2019). Prior to DMA, seed aliquots from the various CD regimes were all re-equilibrated to the same WC (about 0.04 g H_2_O g^-1^ DW).

After removing the seed coat with a scalpel, the visco-elastic properties of the endosperm of seeds equilibrated at defined RHs were determined with a DMA-1 analyser (Mettler Toledo GmbH, Greifensee, Switzerland) over temperatures ranging from −120 to +90 °C. The chosen seed WCs were in equilibrium with the RHs used for CD and extended from 9 to 85% RH. The DMA tests were conducted in compression mode, using spacers to allow clamping of individual seeds in a 1-mm gap. DMA scans of individual seeds were acquired on at least two different seeds for each WC. Static and dynamic forces were set at 200 and 165 mN, respectively, and delivered at a frequency of 1 Hz (Ballesteros and Walters, 2011, 2019). Prior to analyses, samples were cooled from RT to −120 °C in about 10 min using a stream of liquid nitrogen. Thereafter, samples were held isothermally at −120 °C for 1 min and heated to 90 °C at a rate of 3 °C min^-1^. Storage modulus, loss modulus, and tan δ (i.e. loss modulus/storage modulus) were calculated from the heating scans using the software Stare v12.0 (Mettler Toledo, Greifensee, Switzerland), and only tan δ curves were used to measure the different structural relaxations. Large steps or first order peaks of tan δ are related to structural relaxations and phase changes in the seed cytoplasm, which can indicate either the transition from solid to fluid of amorphous solids or the melting of lipid and water crystals (Ballesteros and Walters, 2011). Peaks of tan δ are conventionally labelled with Greek letters (α, β, γ, etc.) from the highest to the lowest temperature, and α relaxations typically correspond to the largest signal in DMA scans (Ballesteros and Walters, 2011). The Tg was determined from the α relaxation peaks, as previously characterised in other seeds and fern spores (Ballesteros and Walters, 2011, 2019; López-Pozo *et al*., 2019).

#### Differential Scanning Calorimetry

The melting transitions of seed storage lipids (i.e. triacylglycerol [TAG]) were detected and characterised using DSC analyses (Vertucci, 1992; Crane *et al*., 2003; Walters *et al*., 2005b), enabling an extensive comparison of the physical and structural status of *P. densiflora* seeds after CD at different RHs under normoxia and hypoxia. As for DMA, aliquots of seeds subjected to the different CD regimes were equilibrated at the same WC of 0.04 g H_2_O g^-1^ DW. After removing the seed coat and excising embryonic axes, melting transitions were determined on both embryonic axes and endosperm using a differential scanning calorimeter DSC-1 (Mettler-Toledo, Greifensee, Switzerland), calibrated for temperature (156.6 °C) and energy (28.54 J g^-1^) with indium standards. Samples were cooled from 25 to −150 °C at a rate of 10 °C min^-1^, held isothermally for 1 min, before heating from −150 to 90 °C at a rate of 10 °C min^-1^. TAG melting transitions were detected as first order transitions (i.e. peaks) from seed heating thermograms (Vertucci, 1992; Crane *et al*., 2003; Walters *et al*., 2005b; Ballesteros and Walters, 2007b). The onset temperature of the TAG melting transitions was calculated from the intersection between the baseline and a line drawn from the steepest portion of the transition peak. Multiple peaks were detected for the TAG melting transitions and represented diverse TAGs or diverse crystalline structures of the same TAG, depending on their melting temperature (Crane *et al*., 2003; Walters *et al*., 2005b; Ballesteros and Walters, 2007b). The enthalpy (ΔH) of the total TAG melting transition was obtained from the area encompassed by all lipid peaks (i.e. L1 and L2) and the baseline (Ballesteros and Walters, 2007b). All analyses were performed using Mettler-Toledo Stare software version 12.0 (Mettler-Toledo, Greifensee, Switzerland). Scans were initially acquired using separated embryos and endosperm, indicating that the melting of TAGs was equivalent in both seed structures (data not shown). However, all results from the DSC analyses presented in this paper refer to seed endosperm only, because six to ten embryonic axes per replicate were required to obtain sufficient signal in the DSC scans, compared to the endosperm of individual seeds. Enthalpies of exothermic and endothermic events were expressed on a DW basis, after drying seed endosperms to 0.04-0.05 g H_2_O g^-1^ DW in chambers set at RT and various RHs as described in (Ballesteros and Walters, 2007b). For each CD regime, DSC scans were acquired on at least four seed endosperms, used as replicates.

#### Water sorption isotherms

Water sorption isotherms were constructed at 45 °C (i.e. the temperature used for the CD regimes under normoxia and hypoxia) for RHs ranging between 0.5 and 75%. WC-RH data for RH ≤ 40% were fit to the BET model to calculate parameters related to surface area and chemical affinity for water or frozen-in structure of glasses, as described earlier for seeds and fern spores (Ballesteros and Walters, 2007b, 2011, 2019). After recording the FWs, seeds were dried at 103 °C for 16 h to obtain the DWs. Seed WCs were calculated as the difference between FW and DW and expressed as g H_2_O g^-1^ DW. For each RH, the WCs of five individual seeds were determined between 7 and 30 d after incubation in RH chambers (i.e. the period during which WC reached a steady-state) and averaged.

### Biochemical analyses

After CD, pools of 40 seeds for each CD regime and replicate, including control seeds from 2015 and seed bank seeds, were immediately frozen in liquid nitrogen and lyophilised for 5 d. Seed WC was expressed as g H_2_O g^-1^ DW after recording FW (i.e. before lyophilisation) and DW (i.e. after lyophilisation) with an XS105 analytical balance (Mettler Toledo GmbH, Columbus, OH, USA). Material for analyses was obtained from seeds pre-cooled for 15 min in 5-mL Teflon capsules (Sartorius GmbH, Göttingen, Germany), containing a 10 mm-diameter agate bead, and ground to a fine powder using a Mikro-Dismembrator S (B. Braun, Biotech International, Melsungen, Germany) at 3,000 rpm for 30 s. Until analysis, ground samples were stored at −80 °C in a hermetically sealed plastic container with silica gel. All biochemical analyses were conducted using ground seed powder and all chemicals listed hereafter were of analytical grade and purchased from Sigma-Aldrich (St. Louis, MO, USA), unless otherwise specified. All solutions were prepared in UPW.

#### HPLC analysis of low-molecular-weight thiol-disulphide redox couples

For each replicate (*n* = 4), 50.0 ± 0.6 mg of seed powder was combined with 24.9 ± 0.8 mg of polyvinylpolypyrrolidone, and thiols and disulphides were extracted in 1 mL of ice-cold 0.1 M HCl using a Tissue-Lyser (Qiagen, Hilden, Germany) and two 3-mm glass beads (30 Hz, 4 min). After a first centrifugation step (28,000 g, 20 min, 4 °C), 700 μL of the supernatants was promptly transferred to a new Eppendorf tube and further centrifuged (28,000 g, 20 min, 4 °C), according to (Schausberger *et al*., 2019). Thereafter, extracts were divided into two separate aliquots: 120 μL for the quantification of both LMW thiols and disulphides (aliquot A), and 400 μL for the quantification of disulphides only (aliquot B). Briefly, after verifying that the pH of extracts lay between 8.00 and 8.30, dithiothreitol (DTT) was used to reduce disulphides in aliquot A. To determine disulphides only, thiols of aliquot B were first blocked with *N*-ethylmaleimide before reduction by DTT. In both aliquots, thiols were derivatised with monobromobimane for detection by fluorescence (excitation: 380 nm; emission: 480 nm) after separation of cysteine (Cys), γ-glutamyl-cysteine (γ-Glu-Cys), cysteinyl-glycine (Cys-Gly), and GSH, using a reserved phase HPLC 1100 system (Agilent Technologies, Inc., Santa Clara, CA, USA) with a ChromBudget 120-5-C18 column (250 x 4.6 mm, 5.0 μm particle size, Bischoff GmbH, Leonberg, Germany). The concentrations of LMW thiols and corresponding disulphides were calculated using external standards and by subtracting the concentration of disulphides (in thiol equivalents) from the concentration of thiols and disulphides, as described earlier (Bailly and Kranner, 2011).

#### Calculation of E_GSSG/2GSH_

The glutathione half-cell reduction potential (E_GSSG/2GSH_) was calculated from the molar concentrations of GSH and GSSG, estimated using seed WCs (expressed as g H_2_O g^-1^ seed DW), according to the Nernst equation (equation 2):

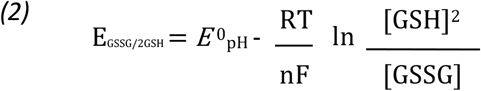

where *R* is the gas constant (8.314 J K^-1^ mol^-1^); *T*, temperature in K; *n*, number of transferred electrons (2 GSH → GSSG + 2 H^+^ + 2 e^-^); F, Faraday constant (9.649 x 10^4^ C mol^-1^); *E*^0^_pH_, standard half-cell reduction potential (*E*^0^’) of a thiol-disulphide redox couple at a defined pH (Schafer and Buettner, 2001; Kranner *et al*., 2006).

In thiol-disulphide redox couples, the concentration of hydrogen ions affects the half-cell reduction potential (E_hc_) (Wardman, 1989), therefore the cytoplasmic pH of control, CD-aged, and seed bank seeds was estimated as previously reported by (Nagel *et al*., 2019) with minor modifications. For each treatment, four replicates of 50.23 ± 0.52 mg of ground seed powder were suspended in 1.2 mL of UPW and shaken at 600 rpm and 100 °C for 10 min. Following centrifugation (15,000 g, 30 min, RT), the supernatants were transferred to fresh Eppendorf tubes and their pH measured using a Multi 3410 pH meter with an ADA S7MDS electrode (VWR International, Wien, Austria). To account for acidification due to interfering compounds released from organelles during extraction of seed powder, a correction factor of +0.6, obtained as difference between the cellular physiological pH (7.30) and the highest pH measured in extracts of control seeds (6.70), was applied as detailed by (Nagel *et al*., 2019). The *E*^0^_pH_ was calculated using the average cytoplasmic pH of each extract according to equation 3:

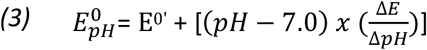

where *E^0’^* is the standard half-cell reduction potential of a thiol-disulphide redox couple at an assumed cellular pH of 7.0 (*E*^0’^_GSSG/2GSH_ = −258 mV), and ΔE/ΔpH refers to the change in the E_hci_ in response to a one-unit pH change. This value equals −59.1 mV at 25 °C for all LMW thiols (Schafer and Buettner, 2001). To show the effect of CD on E_GSSG/2GSH_ without the influence of different seed WCs, the E_GSSG/2GSH_ values of seeds before CD were also estimated at each WC corresponding to the four RHs used for CD.

#### HPLC analysis of tocochromanols

Tocochromanols in 50.3 ± 0.4 mg DW of ground seed powder were extracted in 750 μL of icecold heptane, using two 3-mm diameter glass beads (Carl Roth GmbH+Co, Karlsruhe, Germany) and a Tissue-Lyser (Qiagen, Hilden, Germany) at 25 Hz for 2 min. After centrifugation (28,000 g, 40 min, 4 °C), tocochromanols in 20 μL of supernatant were separated by an HPLC 1100 system (Agilent Technologies, Inc., Santa Clara, CA, USA) on a LiChroCART^®^ column (LiChrospher 100 RP-18, 125 x 4 mm, 5.0 μm particle size, Merck KGaA, Darmstadt, Germany), with constant flow rate of 1 mL min^-1^ of 100% solvent A (acetonitrile: methanol = 74:6) from 0 to 4 min, followed by a gradient changing with linearity to 100% solvent B (methanol: hexane = 5:1) between 4 and 9 min and maintained at 100% up to 20 min. Tocochromanols were detected by fluorescence (excitation: 295 nm; emission: 325 nm) and identification and quantification were based on authentic external standards of α and γ-tocopherol.

#### uHPLC-MS/MS analysis of aldehydes and RES

LMW carbonyls in 51.58 ± 2.17 mg DW of ground seed powder were extracted in 1 mL of acetonitrile containing 0.5 μM 2-ethylhexanal (as internal standard) and 0.05% (w/v) butylated hydroxytoluene, by shaking with two 3-mm glass beads for 2 min at 30 Hz with a Tissue-Lyser (Qiagen, Hilden, Germany). After 5 min in an ice-cold ultrasonic bath, extracts were incubated at 60 °C for 30 min before centrifugation (21,500 g, 20 min, 4 °C). The supernatant was removed, and 12.5 μL of 20 mM 2,4-dinitrophenylhydrazine (DNPH) dissolved in acetonitrile and 19.4 μL of formic acid were added to the pellet and incubated at RT for 1 h in the dark. Before injection, samples were diluted 50:50 with UPW. LMW carbonyls were separated using a reversed-phase column (NUCLEODUR C18 Pyramid, EC 50/2, 50×2 mm, 1.8 μm, Macherey-Nagel, Düren, Germany), using an Ekspert ultraLC 100 UHPLC system (AB SCIEX, Framingham, MA, USA) coupled to a QTRAP 4500 MS for quantification of DNPH-derived aldehydes, according to (Roach *et al*., 2017). Selected carbonyl-DNPH compounds were also quantified using external standards, which were processed and derivatised as for samples and are shown in Supplementary Fig. S4. Peak areas were normalised relative to the internal standard and concentrations were calculated according to calibration curves using the software Analyst and MultiQuant (AB SCIEX, Framingham, MA, USA).

#### Seed oil content, electrical conductivity, and GC-MS analysis of fatty acids

*P. densiflora* seeds were non-invasively quantified for their total oil content using time-domain nuclear magnetic resonance (TD-NMR), according to (Castillo-Lorenzo *et al*., 2019). Three replicates of 15 − 20 intact seeds, equilibrated to ~30% RH, were placed in a Bruker mq20 minispec (Bruker, Coventry, UK) with a 0.47 Tesla magnet (20 MHz proton resonance frequency) at 40 °C, using a 10-mm probe assembly (H_2_o-10-25AVGX4). The method acquired 16 scans with a recycle delay of 2 s. Quantification was achieved by using sunflower oil for calibration, and data were expressed as percentage of oil content (w/w).

Electrolyte leakage during imbibition was used as indicator of membrane integrity (Matthews and Powell, 2006). Control, CD-aged, and seed bank seeds were rinsed with UPW for 15 s to remove surface-bound particles, before imbibing in 6 mL of UPW equilibrated at 20 ± 0.5 °C. During sample stirring at this constant temperature, the electrical conductivity (EC) of leachates released from 25 seeds was measured with a Cond 330i conductivity meter (WTW Xylem Analytics Germany Sales GmbH & Co. KG, Weilheim, Germany) connected to a TetraCon^®^ 325 measuring cell probe, 4 h after the onset of seed imbibition. The values were normalised to seed DW, after drying samples at 103 °C for 17 h.

FAs were quantified after derivatisation to FA methyl esters (FAMEs) via GC-MS, as described by (Li-Beisson, 2010). The transesterification reaction was initiated by mixing 10.14 ± 0.40 mg of finely ground and freeze-dried seed powder in 2 mL of a mixture of methanol: toluene: sulphuric acid (10:3:0.25, v:v:v) supplemented with 0.01% (w/v) butylated hydroxytoluene and containing 200 μg of heptadecanoic acid (solved in methanol: toluene, 10:3, v/v) as internal standard. After incubation at 80 °C and 600 rpm for 90 min, samples were cooled down to RT, before adding 760 μL of hexane and 2.3 mL of 0.9% (w/v) NaCl. Thereafter, samples were vortexed at full speed and centrifuged (3,000 g, 10 min, RT). The supernatants were collected in autosampler vials, injected and FAMEs separated using a Trace 1300 GC coupled to a TSQ8000 triple quadrupole MS (Thermo-Scientific, Waltham, MA, U.S.A.), equipped with a 30-m Rxi-5Sil MS column including a 10-m integra-guard pre-column (Restek Corporation, Bellefonte, PA, USA). A commercial FAMEs mix (Sigma Aldrich ref. 18919, Missouri USA) was used to confirm the identity of the FAMEs. Data analysis was performed using the Xcalibur software v. 4.2 (Thermo-Scientific, Waltham, Massachusetts, USA).

### Statistics

All data were assessed for significance at α = 0.05 using the SPSS Statistics software package v. 25 (IBM, New York, NY, USA). CD under normoxia at low RH (i.e. 9 and 33%) resulted in different seed viability compared to hypoxia, thus individual *t*-tests were run to compare control seeds before CD with seeds exposed to each individual CD regime. Additional *t*-tests were run to compare the effects of O_2_ on biochemical and biophysical measurements between seeds aged at the same RH. The assumption of normal distribution was verified via Shapiro-Wilk test and analysis of quantile-quantile plots. Total germination (%) and WC (% FW) values were arcsine transformed to simulate normal distribution. The assumption of homoscedasticity of variances was assessed through Levene’s test and analysis of the residuals plotted against fitted values. Whenever the latter assumption was not fulfilled, Box-Cox transformations (e.g. log, square root, reciprocal) were applied to the data before analysis. In each dataset, the cut-off value for the Cook’s distance was set at 4/*n* (where *n* was the number of observations in a certain dataset), and all values with a Cook’s distance greater than 4/*n* were considered as outliers and disregarded. Provided that the residuals were not normally distributed, bias-corrected accelerated bootstrap analyses were run with a sample size of 10^5^ and two different seeds (i.e. 2000 and 200), using a Mersenne Twister random number generator algorithm. The 95% confidence intervals generated by bootstrap analyses showed seed sensitivity at the decimal digit.

## Results

### The cytoplasm was glassy at 9% and 33% RH and fluid at 64% and 85% RH, whereas storage lipids always remained fluid during CD at 45 °C

The physical properties of *P. densiflora* seeds at the moisture conditions used in all CD regimes were assessed combining information from DMA, DSC, and water sorption isotherms. In the DMA scans, α relaxations denoted the temperature at which the amorphous solid structure of seed cytoplasm (i.e. the glass) melted into a fluid system, which is indicative of the Tg. Similar to the Tg, α relaxation in non-aged control seeds shifted towards lower temperatures as the seed WC increased (Fig. 1A). Notably, the temperature and size of the α relaxations measured by DMA, or the Tgs detected as second order transitions by DSC, were not significantly affected by the CD regimes (data not shown). The DMA scans also revealed two further structural relaxations below the water freezing point. These structural relaxations were not affected by seed WC and were attributable to melting events of the FAs of seed storage lipids, particularly TAGs. The highest and sharpest peak (named L1 instead of β relaxation to avoid confusion with the β relaxations occurring within the aqueous matrix) appeared between ~-100 and −80 °C, followed by a second less prominent and broader one (L2), extending from ~-80 °C to ~-20 °C (Fig. 1A). The presence of lipid peaks in the endosperm was consistent with a high seed oil content of 29.7 ± 1.2% (w/w) on a fresh weight (FW) basis, quantified with TD-NMR and also revealed by DSC. Furthermore, DSC analyses targeting the hydrophobic domain of seed endosperm clearly detected melting peaks of storage lipids in the same temperature range of L1 and L2 (Supplementary Fig. S1), confirming the lipid nature of these two relaxations.

**Fig. 1.**
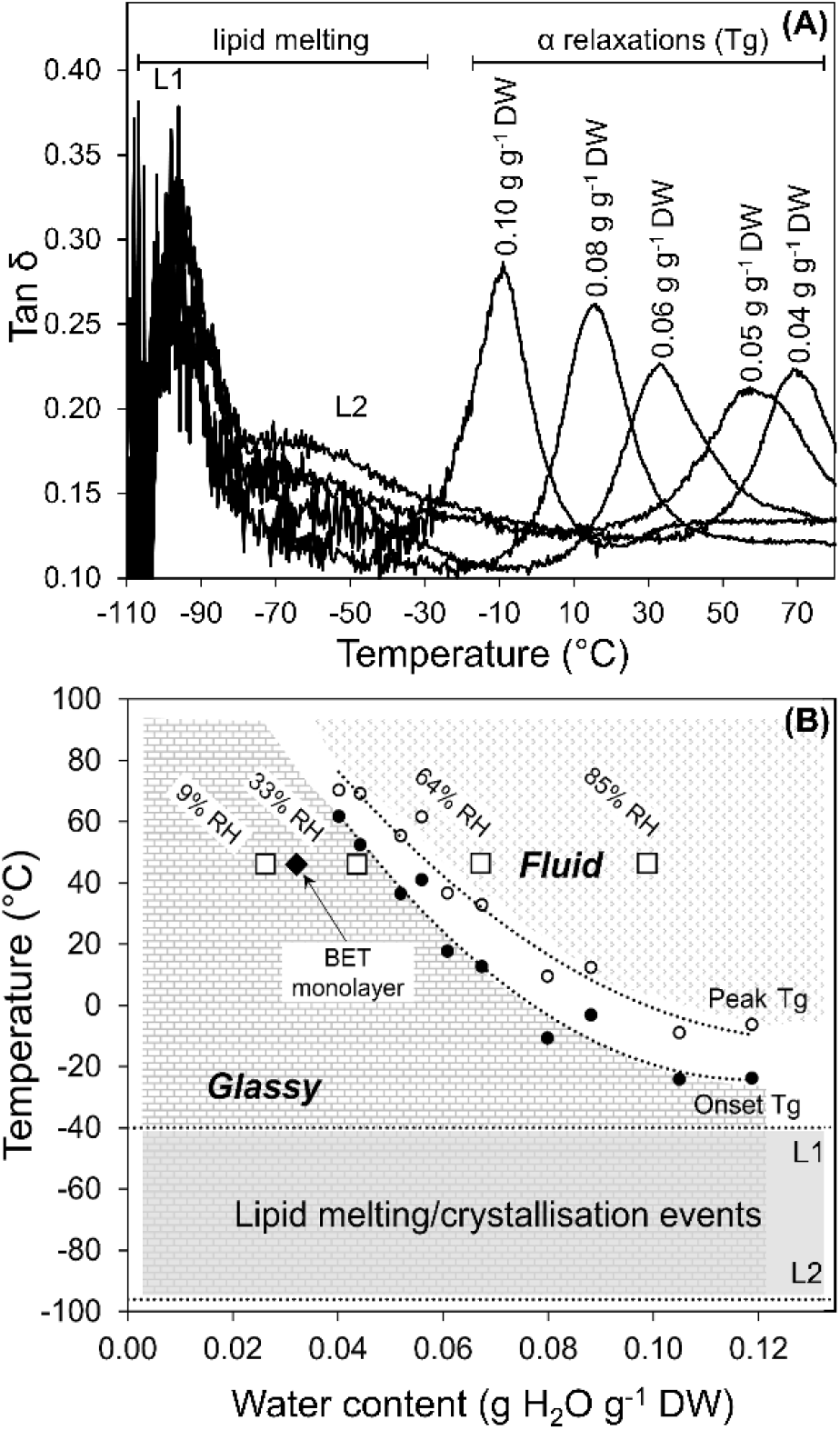
Physical state of *Pinus densiflora* seeds in relation to water content (WC) and temperature. (A) Representative dynamic mechanical analyses (DMA) scans of seeds before controlled deterioration at diverse WCs, expressed as g H_2_O per g dry weight (g g^-1^ DW). Scans show the tan δ, which is the coefficient between the loss modulus and the storage modulus and measures the damping function related to molecular mobility. Peaks in tan δ are indicative of diverse types of structural relaxations. The glass transition temperature (Tg) is characterised by the α relaxation peak, which moves to lower temperatures as the sample WC increases. The melting of storage lipids is indicated by two peaks (L1 and L2), which occur within the same temperature range independently of the sample WC. L1 and L2 were further characterised by differential scanning calorimetry (DSC; refer to Supplementary Fig. S4). (B) Phase diagram constructed using DMA and DSC data. The terms “glassy” and “fluid” refer to the aqueous domain of the cytoplasm (measured by DMA), while L1 and L2 correspond to the melting peaks determined by DSC and define the range of WCs and temperature, at which the hydrophobic domain (i.e. seed storage lipids) showed physical changes. The Tg is depicted by the area within the onset (closed circles) and the peak (open circles) of the α relaxations measured by DMA. The diamond indicates the Brunauer-Emmet-Teller (BET) monolayer, calculated from water sorption isotherms of seeds equilibrated at 45 °C. White squares denote the seed WCs reached at 45 °C at the various relative humidities (RH; n ≥ 2 seeds for each WC).

Based on DMA and DSC analyses, in seeds aged at 45 °C the transition from glassy to fluid cytoplasm started at a seed WC of 0.05 g H_2_O g^-1^ DW, reaching a peak at 0.06 g H_2_O g^-1^ DW, which corresponded to RHs of 42 and 50%, respectively, as per the water sorption isotherms (Fig. 1B). Therefore, the aqueous phase of the cytoplasm of seeds treated at 9 and 33% RH (corresponding to 0.027 and 0.042 g H_2_O g^-1^ DW, respectively) was in a glassy state with restricted molecular mobility (Fig. 1B). In contrast, seeds exposed to CD at 64 and 85% RH (corresponding to 0.069 and 0.098 g H_2_O g^-1^ DW, respectively) had WCs above the Tg and were aged with a fluid cytoplasm and higher molecular mobility (Fig. 1B). Water sorption isotherms at 45 °C enabled to calculate the BET monolayer, which corresponded to a seed WC of 0.033 g H_2_O g^-1^ DW or 18% RH (Fig. 1B). Knowledge of the BET monolayer value contributed to further characterise the glassy state, indicating that during CD at 9% RH not all water binding sites of the surface of macromolecules were saturated (i.e. the BET monolayer was not complete, as from the BET adsorption model). However, during CD at 33% RH, all water binding sites of macromolecules became occupied by water molecules, forming a complete BET monolayer. Furthermore, DMA and DSC analyses showed that the seed storage lipids remained fluid during all the diverse CD regimes at 45 °C (Fig. 1B). Finally, the physical properties of non-aged control seeds suggested that seed bank seeds with a WC of 0.06 ± 0.01 g H_2_O g^-1^ DW (determined after lyophilisation) were in the glassy state during storage at 4 and −20 °C. Based on the cooling and heating DSC scans (Supplementary Fig. S1; cooling scans not shown), seed storage lipids seeds were crystallised during storage at −20 °C, fluid during storage at 4 °C, and completely thawed when seeds had germinated at 20 °C.

### Hypoxia prevented loss of viability only when seeds were aged in the glassy state

In control seeds before CD, total germination was 90%, and seeds required about 12 days to reach the T_25_, here used as an indicator of germination rate. After CD under normoxia, seed viability was significantly impaired, as indicated by lower total germination, longer T_25_, and enhanced electrolyte leakage during initial imbibition. The response to O_2_ concentrations differed depending on the seed physical state (Fig. 2). Overall, seeds exposed to CD died faster at higher RHs (Fig. 2; Supplementary Figs. S2, S3; Supplementary Table S1). At low RH (i.e. in the glassy state), CD resulted in significantly decreased seed viability more under normoxia than hypoxia. For instance, after 138 days of CD at 9% RH, seeds aged under normoxia did not germinate, whereas seeds under hypoxia retained total germination and germination rate (~12 d) comparable to the non-aged control (*P*-value > 0.05; Fig. 2A, B, Supplementary Fig. S2). Similarly, after 70 days of CD at 33% RH, ageing under hypoxia resulted in 2.3-fold higher total germination and faster germination rate compared to normoxia (Fig. 2A, B). The deleterious effects of O_2_ on the viability of glassy-state seeds were also revealed by significantly increased electrolyte leakage from seeds aged under normoxia, which was about 3- and 2-fold higher at 9 and 33% RH, respectively, compared to seeds aged under hypoxia (Fig. 2C). In contrast, during CD seeds with fluid cytoplasm (i.e. at 64 and 85% RH) reached comparable total germination under both normoxia and hypoxia (on average 66%), had similar germination rates (16-17 d), and electrolyte leakage did not significantly differ (Fig. 2, Supplementary Fig. S3).

**Fig. 2.**
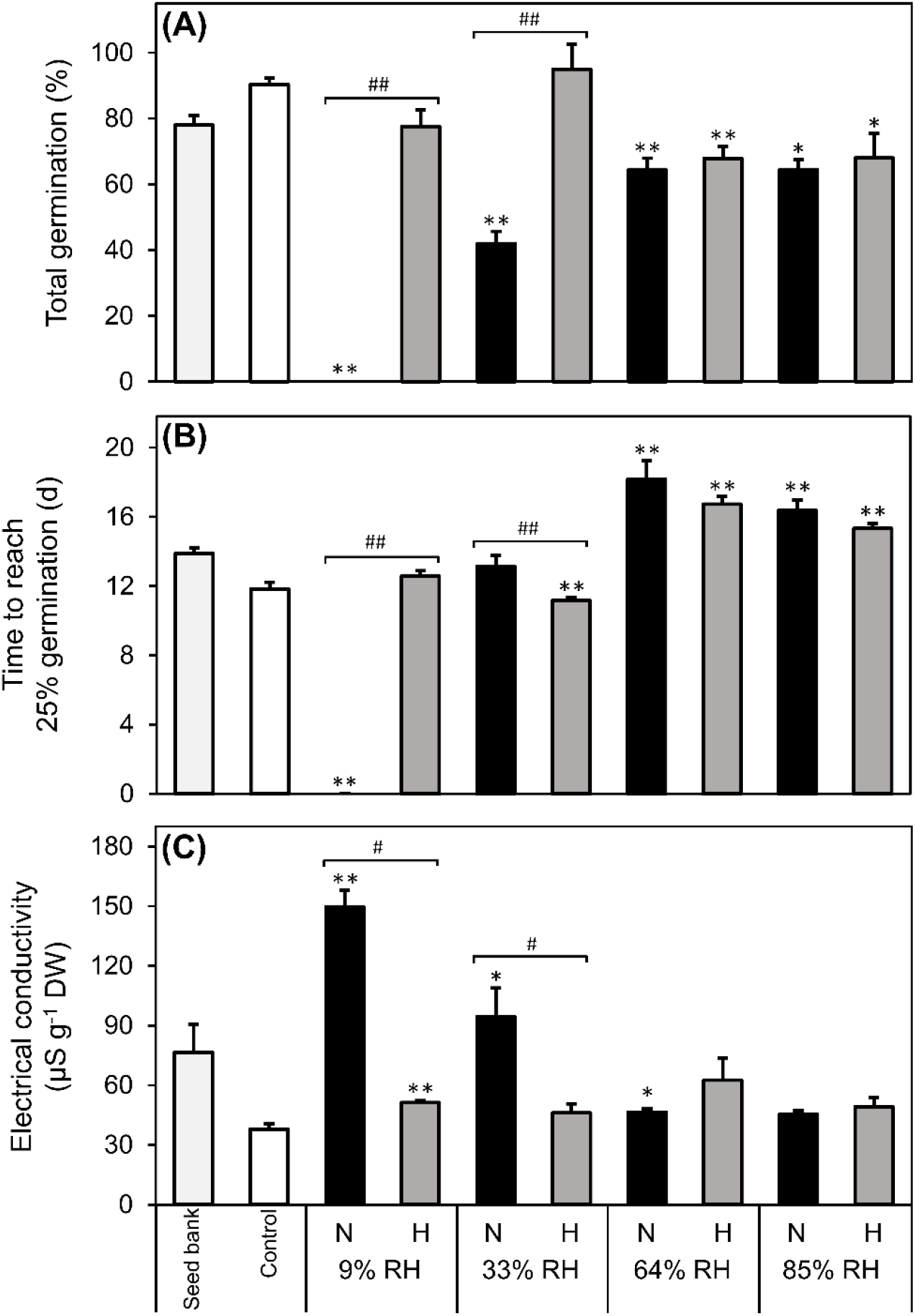
Effects of relative humidity (RH) on the influence of O_2_ during controlled deterioration (CD) at 45 °C on germination and electrolyte leakage. *Pinus densiflora* seeds were aged under normoxia (N, black bars, 19.6% O_2_) and hypoxia (H, grey bars, 0.4% O_2_) at indicated RHs. (A) Total germination (TG) measured after 45 days. (B) Time to reach 25% TG. Data were calculated from the germination curves shown in Supplementary Fig. S2. (C) Electrical conductivity of seed leachates. Asterisks denote significant differences (*, *P*-value < 0.05; **, *P*-value < 0.01) after *t*-tests comparing seeds before CD (control, white bars) with seeds exposed to CD at four RHs under N or H. Hash symbols denote significant differences (#, *P*-value < 0.05; ##, *P*-value < 0.01) after *t*-tests between seeds aged under N and H at the same RH. Values of seeds stored for 20 years at ~0.06 g H_2_O g^-1^ dry weight and low temperatures (seed bank, light grey bars) are also shown. Data are means (*n* = 4 replicates of 50 seeds each) ± SE.

Notably, seeds aged at 9% RH under normoxia died faster than predicted using the regression obtained from P50 values at higher RHs (Supplementary Fig. S2; Supplementary Table 1). Finally, longterm cold storage of seed bank seeds resulted in significantly lower total germination (78%, *P*-value < 0.01) compared to initial viability after harvest (91%; data not shown), and the electrolyte leakage from seed bank seeds was about twice than from control seeds (Fig. 2C).

### GSH concentrations declined during ageing and independently of O_2_ in seeds with a fluid cytoplasm

The water-soluble antioxidant GSH was the most abundant LMW thiol (cf. Fig. 3A and Supplementary Fig. S4), and CD led to a conversion of GSH to GSSG (Fig. 3A). In the glassy state (i.e. CD at 9 and 33% RH), normoxia led to a > 50% drop in GSH concentrations, whereas under hypoxia GSH declined by only 12%. This agreed with seeds accumulating 1.3 to 1.5-fold more GSSG under normoxia than hypoxia at 9 and 33% RH, respectively (*P*-values = 0.004 and 0.001; Fig. 3A). Ageing seeds with fluid cytoplasm (i.e. CD at 64 and 85% RH) led to an 80% drop in GSH concentrations, and at 85% RH under normoxia significantly more GSH was consumed and GSSG accumulated than under hypoxia (*P*-values < 0.001 and 0.004, respectively; Fig. 3A). Consequently, the oxidative shift in E_GSSG/2GSH_ was larger in seeds aged under normoxia than under hypoxia after CD at 9, 33, and 85% RH (Fig. 3B). Of note, GSH decreases prevailed over GSSG accumulation in seeds with fluid cytoplasm during CD, leading to > 40% loss of total glutathione (i.e. GSH + GSSG) when calculated as GSH equivalents (GSSG = 2 GSH). Seed WC was used to estimate the molar concentrations of GSH and GSSG, and GSH is a squared term in the Nernst equation to calculate E_GSSG/2GSH_ (equation 2). Therefore, seeds with different WCs, but with the same GSH and GSSG molar concentrations, will have different E_GSSG/2GSH_ values on a DW basis (note differences between the open circles in Fig. 3B, indicating respective E_GSSG/2GSH_ values of seeds at each WC in equilibrium with chosen RHs before CD). At 9% RH, seed WC was just 0.4% FW, and after CD a net increase in GSH molar concentrations occurred relative to control seeds (WC = 3.9% FW), despite GSH consumption on a DW basis (Fig. 3A). Conversely, at 85% RH a higher seed WC of 7.9% FW diluted GSH, resulting in E_GSSG/2GSH_ less negative values (i.e. more oxidising conditions; Fig. 3B). Nonetheless, GSSG accumulation and mainly GSH consumption were major factors contributing to the oxidative shift of E_GSSG/2GSH_ in seeds aged with fluid cytoplasm (Fig. 3B). In seed bank seeds, GSH concentrations dropped by 42% in comparison to control seeds, leading to less negative values of E_GSSG/2GSH_ (Fig. 3A, B).

**Fig. 3.**
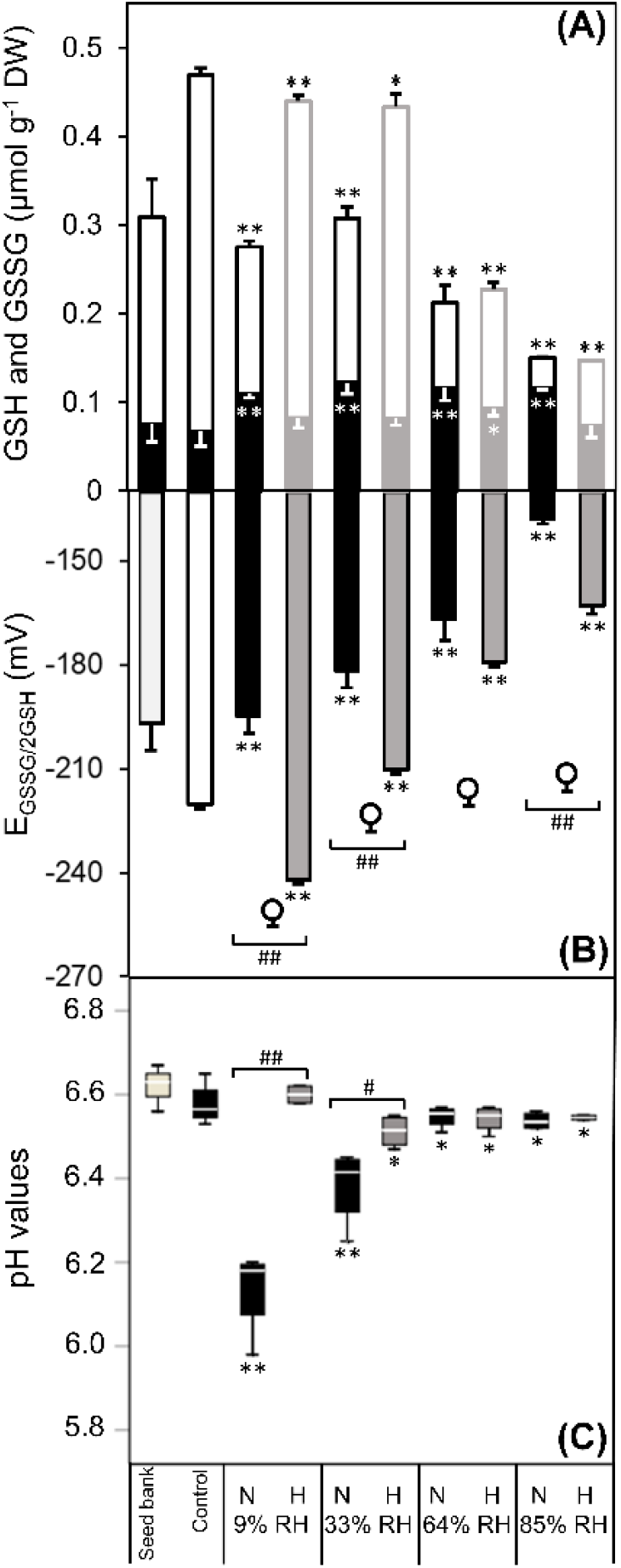
Effects of relative humidity (RH) on the influence of O_2_ during controlled deterioration (CD) at 45 °C on glutathione concentrations and redox state, and pH of seed extracts. *Pinus densiflora* seeds were aged under normoxia (N, black bars, 19.6% O_2_) and hypoxia (H, grey bars, 0.4% O_2_) at indicated RHs. (A) Concentrations on a dry weight (DW) basis of the low-molecular-weight thiol glutathione (GSH, open bars) and its disulphide (GSSG, closed bars). (B) Half-cell reduction potential of the GSSG/2GSH redox couple (E_GSSG/2GSH_) calculated according to the Nernst equation at the pH values showed in panel (C) with an offset correction. Circles indicate the E_GSSG/2GSH_ values calculated using the seed water contents at the end of seed pre-equilibration at indicated RHs and prior to CD. (C) pH of extracts obtained from finely ground seed powder. Asterisks denote significant differences (*, *P*-value < 0.05; **, *P*-value < 0.01) after *t*-tests comparing seeds before CD (control, white bars and open circles) with seeds exposed to CD at four RHs under N or H. Hash symbols denote significant differences (#, *P*-value < 0.05; ##, *P*-value < 0.01) after *t*-tests between seeds aged under N and H at the same RH. Values of seeds stored for 20 years at ~0.06 g H_2_O g^-1^ DW and low temperatures (seed bank, light grey bars) are also shown. Data are means (*n* = 4 replicates of 50 seeds each) ± SE.

The Nernst equation to calculate E_GSSG/2GSH_ is also dependent on cellular pH values (equation 2). All CD treatments, except for 9% RH under hypoxia, resulted in a significant cellular acidification (Fig. 3C), consequently contributing to more oxidising conditions (a difference in pH of 0.1 influences the E_GSSG/2GSH_ by 6 mV). In general, seed cellular acidification reflected the changes in total germination, whereby loss of seed viability was accompanied by lower pH (Figs 1A, 3C). The pH of seeds aged with a glassy cytoplasm decreased only marginally under hypoxia, whereas seeds aged with a fluid cytoplasm showed a slight but significant acidification regardless of O_2_ concentrations during CD (Fig. 3C). Other LMW thiols included Cys, γ-Glu-Cys, and Cys-Gly. The GSH intermediates total Cys (i.e. Cys + cystine) and total γ-Glu-Cys (i.e. γ-Glu-Cys and bis-γ-glutamyl-cystine) were always more abundant than total Cys-Gly (i.e. Cys-Gly + cystinyl-bis-glycine) (Supplementary Fig. S4). Notably, in the fluid state at 64 and 85% RH, seeds contained on average more total γ-Glu-Cys (2.8-fold) and total Cys (1.9-fold) than the control (Supplementary Fig. S4).

### Unsaturated fatty acids depleted in glassy-state seeds aged under normoxia

DSC analyses enabled to quantify the effects of CD on physical changes of seed storage lipids (mainly TAGs). Melting of seed TAGs was detected as first order peaks in the DSC heating scans (Supplementary Fig. S1). In non-aged control seeds two distinct melting peaks occurred at −96 ± 2 °C (L1) and −40 ± 2 °C (L2), with a total ΔH of lipid melt of 17.9 ± 5.8 mJ g^-1^ DW (Fig. 4A). Seeds deteriorated at various RHs under normoxia and hypoxia also displayed lipid melting peaks between −100 and −70 °C (L1) and between −50 and −5 °C (L2; Supplementary Fig. S1). The onset and peak temperatures of the melting transitions associated to both lipid peaks were not significantly affected by the CD regimes (Supplementary Fig. S1). However, the ΔH of lipid melt was altered by the CD regimes, and significant changes were detected only in seeds aged under normoxia in the glassy state (Fig. 4A), whereby the total ΔH of lipid melt significantly dropped by 3- and 1.5-fold in seeds aged at 9 and 33% RH, respectively (Fig. 4A), and mostly related to peak L1 (Supplementary Fig. S1).

**Fig. 4.**
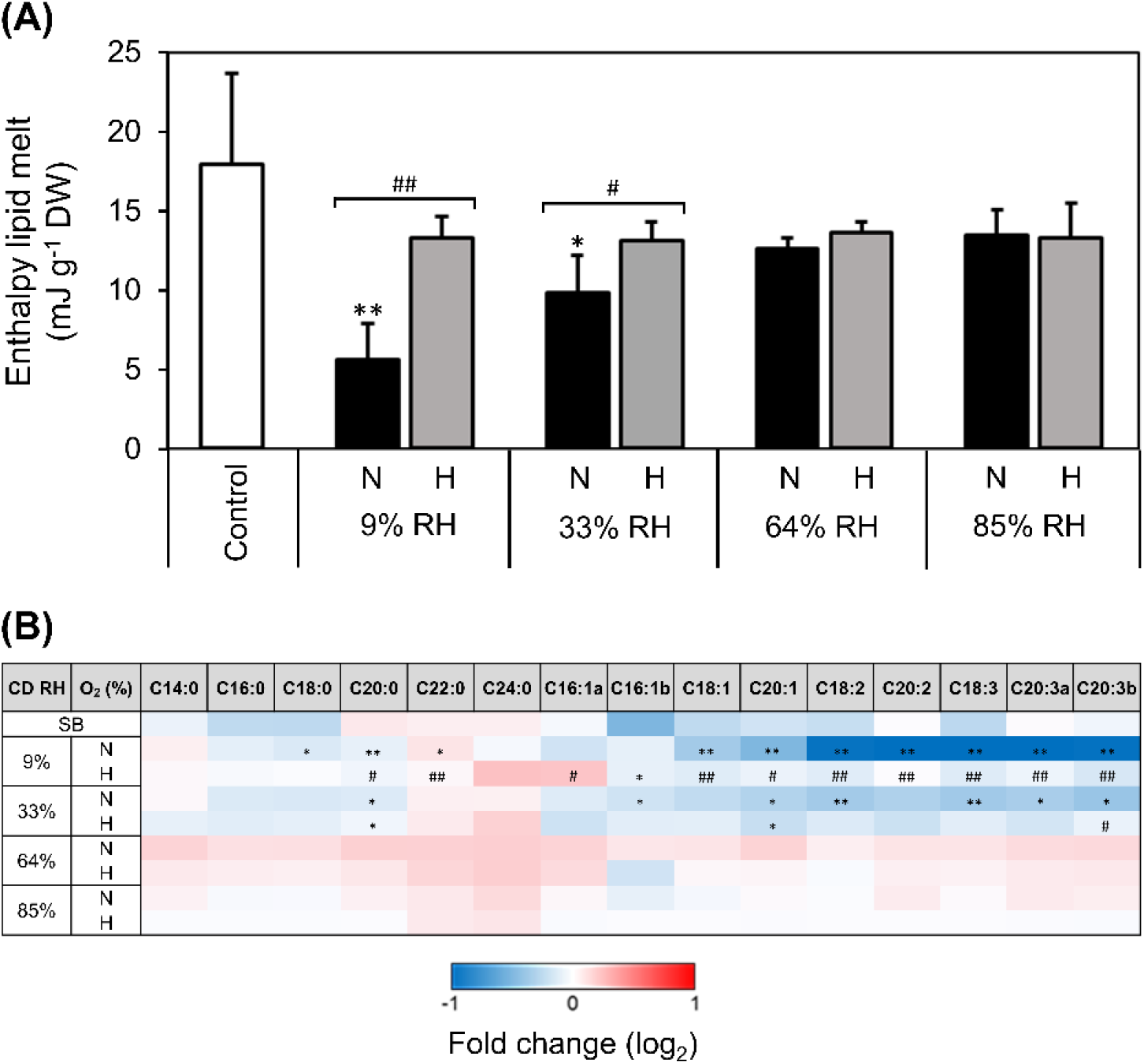
Effects of relative humidity (RH) on the influence of O_2_ during controlled deterioration (CD) at 45 °C on the enthalpy of melt of storage lipids and relative abundance of fatty acids. *Pinus densiflora* seeds were aged under normoxia (N, black bars, 19.6% O_2_) and hypoxia (H, grey bars, 0.4% O_2_) at indicated RHs. (A) Enthalpy of melt of storage lipids calculated from heating scans acquired with differential scanning calorimetry on seed endosperm, after excising embryonic axes. Asterisks denote significant differences (*, *P*-value < 0.05) after *t*-tests between the endosperms of non-aged seeds, used as control, and seeds exposed to CD. Data are means (*n* = 4 seeds) ± SE. (B) Fold-change in the abundance of individual fatty acids, as compared to the non-aged control, measured as fatty acid methyl esters with gas chromatography coupled to mass spectrometry. Differences on a log2 scale are shown by the bottom key highlighting decreases (blue), accumulation (red), and absence of changes (white). Asterisks denote significant differences (*, *P*-value < 0.05; **, *P*-value < 0.01) from *t*-tests comparing seeds before CD (control) with seeds after exposure to CD at four RHs under N or H. Hash symbols denote significant differences (#, *P*-value < 0.05; ##, *P*-value < 0.01) after *t*-tests between seeds aged under N and H at the same RH. Values of seeds stored for 20 years at ~0.06 g H_2_O g^-1^ DW and low temperatures (seed bank, SB) are also shown. Data are means (*n* = 4 replicates of 50 seeds each) ± SE. Different letters (e.g. a and b) refer to fatty acid isomers (with the same number of carbons and double bonds). C14:0, myristic acid; C16:0, palmitic acid; C18:0, stearic acid; C20:0 arachidic acid; C22:0, behenic acid; C24:0, lignoceric acid; C16:1, palmitoleic acid; C18:1 oleic acid; C20:1, eicosenoic acid; C18:2, linoleic acid; C20:2, eicosadienoic acid; C18:3, linolenic acid; C20:3, dihomo-γ-linolenic acid.

To assess if such alterations of seed storage lipids’ physical state were accompanied by chemical changes, the total content of each FA (i.e. constituting membranes and TAG of oil bodies) were measured with GC-MS. The most abundant FAs of *P. densiflora* seeds included linolenic (C_18:3_), palmitic (C_16:0_), linoleic (C_18:2_), oleic (C_18:1_), stearic (C_18:0_), and dihomo-γ-linolenic acid (C_20:3_) (Supplementary Fig. S1). Depletion of FAs with unsaturated carbon bonds, and particularly polyunsaturated fatty acids (PUFAs), occurred in seeds aged under normoxia with a glassy cytoplasm, with hypoxia attenuating these drops (Fig. 4B). In contrast, saturated FAs were much less affected. Notably, no significant changes in any detected FAs occurred in seeds aged with a fluid cytoplasm (Fig. 4B). Seed bank seeds contained less palmitoleic (C_16:1_), oleic (C_18:1_), and linolenic (C_18:3_) acid than control seeds before CD (Fig. 4B).

### Glassy-state seeds aged under normoxia underwent tocochromanols consumption and substantial increases of reactive electrophile species and aldehydes

*P. densiflora* seeds contained about 30-fold more γ-tocopherol than α-tocopherol (Fig. 5). In seeds aged in the glassy state under normoxia, γ-tocopherol concentrations decreased by 8.0 and 2.0-fold at 9 and 33% RH, respectively, and these losses were alleviated under hypoxia. In contrast, γ-tocopherol concentrations did not show pronounced changes after CD in seeds aged with fluid cytoplasm (64 and 85% RH). Additionally, γ-tocopherol concentrations were lower in seed bank seeds compared to control seeds. The much less abundant α-tocopherol was depleted under normoxia at 9% RH, at a seed WC below the BET monolayer value (Fig. 5).

**Fig. 5.**
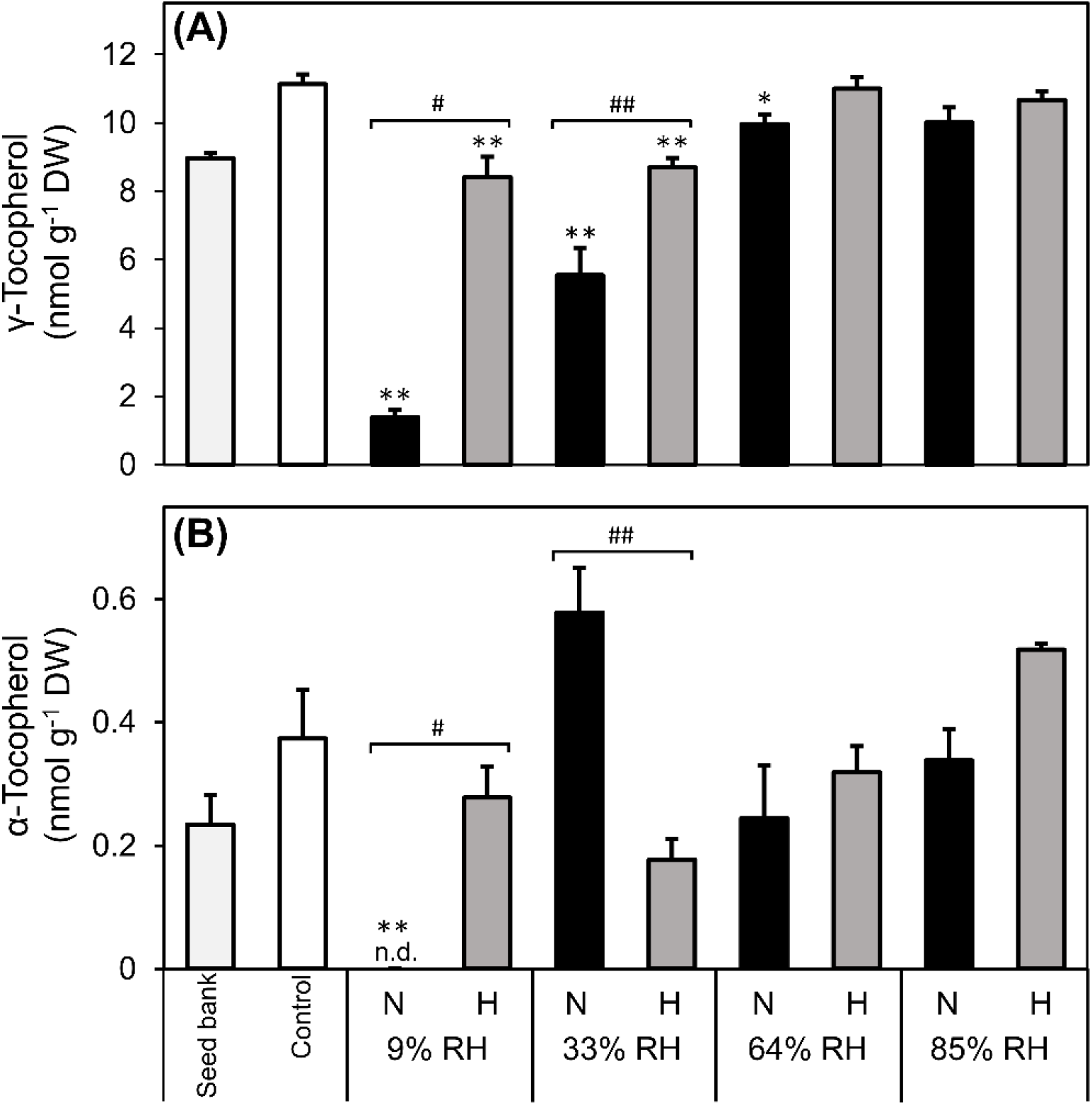
Effects of relative humidity (RH) on the influence of O_2_ during controlled deterioration (CD) at 45 °C on the concentrations of tocopherols. *Pinus densiflora* seeds were aged under normoxia (N, black bars, 19.6% O_2_) and hypoxia (H, grey bars, 0.4% O_2_) at the indicated RHs. Concentrations on a dry weight (DW) basis of (A) γ-tocopherol, and (B) α-tocopherol. Asterisks denote significant differences (*, *P*-value < 0.05; **, *P*-value < 0.01) from *t*-tests comparing seeds before CD (control) with seeds after exposure to CD at four RHs under N or H. Hash symbols denote significant differences (#, *P*-value < 0.05; ##, *P*-value < 0.01) after *t*-tests between seeds aged under N and H at the same RH. Values of seeds stored for 20 years at ~0.06 g H_2_O g^-1^ DW and low temperatures (seed bank, light grey bars) are also shown. Data are means (*n* = 4 replicates of 50 seeds each) ± SE; n.d. = not detected.

Relative to the non-aged control, seeds aged by CD in the glassy state (9 and 33% RH) under normoxia contained more aldehydes, RES, and (di)carboxylic acids (Fig. 6), in agreement with the loss of PUFAs (Fig. 4B). Such increases included > 250-fold more hexanal and azelaic acid, > 50-fold more azelaaldehydic and suberic acids, and > ten-fold more of the RES 4-hydoxynonenal and malondialdehyde. Conversely, seed storage under hypoxia at the same RHs prevented such increments (Fig. 6). Hexanal was by far the most abundant aldehyde detected in aged seeds, either after storage in response to CD or seed bank conditions (Supplementary Fig. S5). Ageing seeds with a fluid cytoplasm resulted in concentrations of acrolein, 4-hydroxyhexenal, trans-2-hexenal, and benzaldehyde falling 2fold below their concentrations in non-aged control, while the accumulation of aldehydes was modest (Figure 6; Supplementary Fig. S5). Notably, these changes were only loosely coupled to O_2_ availability (Fig. 6; Supplementary Fig. S5). Finally, seed bank seeds contained more 4-hydroxynonenal, acrolein, and butyraldehyde than control seeds (Fig. 6; Supplementary Fig. S5).

**Fig. 6.**
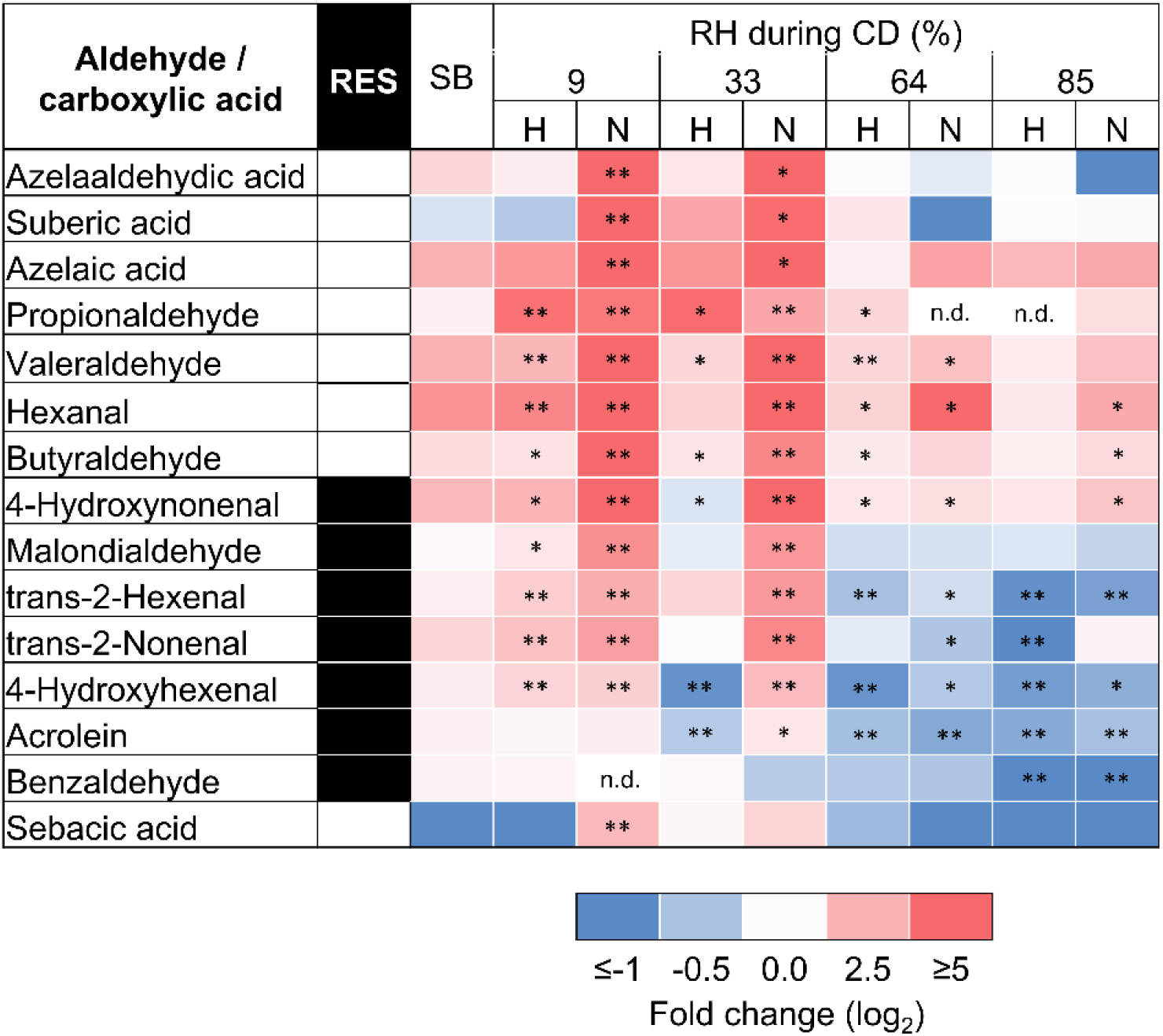
Effects of relative humidity (RH) on the influence of O_2_ during controlled deterioration (CD) at 45 °C on relative amounts of fatty acid breakdown products. *Pinus densiflora* seeds were aged under normoxia (N, 19.6% O_2_) and hypoxia (H, 0.4% O_2_) at indicated RHs. Fold-change in the abundance of aldehydes (measured via ultra-high performance liquid chromatography coupled to mass spectrometry [MS]) and (di)carboxylic acids (measured via gas chromatography coupled to MS) are shown on a log_2_ scale via shading, with blue and red highlighting decline and accumulation, compared to the non-aged control, respectively, as indicated by the key. Pale shading and n.d. indicate no change and when a compound was not detected, respectively. Black boxes next to compound names signify reactive electrophile species (RES). Asterisks denote significant differences (*, *P*-value < 0.05; **, *P*-value < 0.01) from *t*-tests comparing individual species in seeds before CD with seeds after CD at four RHs under H or normoxia N. Values of seeds stored for 20 years with at ~0.06 g H_2_O g^-1^ DW and low temperatures (seed bank, SB) are also shown. Data are means (*n* = 4 replicates of 50 seeds each) ± SE.

## Discussion

Oxygen is directly involved in deteriorative reactions of macromolecules (McDonald, 1999; Bailly, 2004; Kranner *et al*., 2010; Sano *et al*., 2016), but its underlying effect on seed longevity has never been integrated with knowledge on structural mechanics and thermodynamics of seed deterioration. In this paper, we combined biophysical and biochemical analyses of *P. densiflora* seeds to clarify how contrasting physical states within seeds influence the contribution of O_2_ to reactions accompanying ageing.

### The physical state of the cytoplasm determine molecular mobility and affect seed ageing reactions

Seed WC and storage temperature, together with genetic background, hormonal regulation, and environmental conditions experienced during seed development, maturation, and desiccation, all influence orthodox seed longevity (Buitink and Leprince, 2004; Nagel et al., 2015; Leprince et al., 2017; Zinmeister et al., 2020). While genetic background and environmental conditions during seed development establish the biochemical composition of seed cells, seed WC and storage temperature determine the physical state of the cytoplasmic domains, which vary depending on the “dry architecture” of seed cells (Ballesteros *et al*., 2020). This is critical to the longevity of desiccated seeds, because the physical state of aqueous and lipid domains define the physiological events and the rates of physicochemical reactions contributing to seed deterioration (Vertucci and Roos, 1990; Hoekstra *et al*., 2001; Ballesteros *et al*., 2020). Across all CD regimes used in this study, DSC analyses revealed that *P. densiflora* seeds always maintained a liquid lipid domain (e.g. lipid droplets of storage TAGs). However, the seed aqueous domain was in the glassy state when aged by CD at 9 and 33% and became fluid when aged by CD at 64 and 85% RH, as determined by DMA (Fig. 1). Under all CD regimes, the fluid state of the lipid domain would have enabled molecular mobility of the main FA chains and their side groups. However, the activity of cytosolic lipid-metabolising enzymes (e.g. lipases and lipoxygenases that catalyse lipid hydrolysis and oxidation, respectively) would be restricted by the glassy state. In such a highly viscous conditions, molecular mobility is limited to vibration, bending, and rotation of the side groups of macromolecules (Ballesteros and Walters, 2011; Ballesteros *et al*., 2020), which is not sufficient to permit enzymatic catalysis (Fernández-Marín *et al*., 2013; Candotto Carniel *et al*., 2021) but allows diffusion of small molecules, such as O_2_ (reviewed in Ballesteros et al., 2020). In contrast, the molecular mobility of the aqueous matrix of the cytoplasm increased in the fluid state, ensuring the movement of the main chains of macromolecules, which is compatible with enzyme activity (Ballesteros and Walters, 2011). Particularly, enzymes were able to diffuse across the fluid cytoplasm, thus affecting the type of biochemical reactions that lead to seed ageing (Walters, 1998).

Altogether, due to the increased molecular mobility, possibly resuming enzymatic activity in the cytoplasm, seed ageing in the fluid state was accelerated compared to the glassy state (Fig. 2, Supplementary Fig. S3).

### O_2_ is detrimental to the longevity of seeds with a glassy but not fluid cytoplasm

Several studies have shown a detrimental effect of O_2_ on seed longevity (e.g. (Harrison, 1966; Bennici *et al*., 1984; Shrestha *et al*., 1985; Barzali *et al*., 2005; González-Benito *et al*., 2011; Groot *et al*., 2012; Groot *et al*., 2015; Schwember and Bradford, 2011)), in line with a role for ROS in deterioration, as proposed by the “free-radical theory of ageing” (Harman, 1956). However, other studies reported that longevity of seeds aged by CD with a fluid cytoplasm was not influenced by elevated O_2_ (Ohlrogge and Kernan, 1982; Ellis and Hong, 2007; Morscher *et al*., 2015; Roach *et al*., 2018a; Schausberger *et al*., 2019). Here, seeds in the fluid state aged rapidly irrespectively of O_2_ availability (Figs. 1B, 2). (Ibrahim and Roberts, 1983) showed that O_2_ impaired lettuce seed longevity only at WC < 0.18 g H_2_O g^-1^ DW, suggesting that seed WC is a relevant determinant of how O_2_ affects longevity. Altogether, these reports indirectly draw attention to differential ageing mechanisms tied to seed physical state. Particularly, in most of the fore-mentioned studies, in which O_2_ impaired longevity, seeds were likely aged in the glassy state, as estimated according to available temperatures, WCs, and RHs. Our study on *P. densiflora* provides direct evidence that normoxia severely shortened seed longevity only when seeds were in the glassy state (Figs. 1, 2).

Based on a negative logarithmic relationship between seed WC (corresponding to RHs between 30 and 100%) and P50 values under normoxia at 45 °C, a P50 of 248 days for seeds aged at 9% RH was estimated (Supplementary Fig. S2). As such, complete loss of germination of these seeds after only 138 days is indicative of the so-called “critical moisture content” (corresponding with WCs in equilibrium with 10-15% RH at 20 °C), beyond which further decreases in seed WC do not extend longevity (Ellis *et al*., 1990; Ellis *et al*., 1992; Ellis and Hong, 2006). Nonetheless, seeds aged under hypoxia at 9% RH hardly showed any signs of deterioration after 138 d (Fig. 2). Albeit we have insufficient ageing intervals to calculate P50 values under hypoxia, it would take considerably longer to reach the P50 value of glassy-state seeds aged under normoxia at 9% RH. Considering that normoxia did not speed up ageing rates in the fluid state, but that longevity was extended in the glassy state, the negative logarithmic relationship between P50 values and seed WCs would most likely no longer fit under hypoxia, as it did under normoxia (Fig. 2, Supplementary Fig. S2). In a few studies, dehydration below the “critical moisture content” led to more rapid loss of viability than seeds stored with higher WC (Ellis *et al*., 1988, 1989; Vertucci *et al*., 1994). This phenomenon has been related to the removal of the water that is tightly associated with macromolecular surfaces, such as that the BET monolayer on the surface of cytoplasmic macromolecules and lipid droplets (Labuza, 1980; Buitink *et al*., 1998; Ballesteros and Walters, 2007b; Barden and Decker, 2016), which is the physical situation occurring in seeds aged at 9% RH in the present study (Fig. 1B). In seeds dried below the critical moisture content no water is strongly bound to macromolecules, and O_2_ could attack empty water-binding sites of macromolecules, such as oleosins at the surface of lipid droplets and polar residues of lipid bilayers. Oleosins are essential to stabilise the oil bodies of dry seeds during seed imbibition (Leprince *et al*., 1998) and seem to participate to lipid droplet breakdown by recruiting lipases and other hydrolytic enzymes involved in storage lipid metabolism during germination and early seedling growth (Chapman *et al*., 2012). Regardless, the high sensitivity to O_2_ of seeds aged at 33% RH (with a complete BET monolayer) in terms of viability loss, electrolyte leakage, and lipid peroxidation, suggests that the Tg is already a clear WC threshold below which seeds become susceptible to O_2_-mediated deterioration.

### Glutathione conversions and redox state reveal that O_2_ diffusion and ROS production are not totally restricted in the glassy state

To understand the influence of O_2_ on the redox state of the aqueous cytoplasmic domain under contrasting physical states during seed ageing, we focused on the hydrophilic antioxidant GSH. Dry seeds contain much more GSSG than healthy and hydrated plant tissues, and GSH conversion to GSSG is promoted during seed desiccation and ageing (Meyer *et al*., 2007; Colville and Kranner, 2010). Large oxidative shifts of the cellular redox environment, as viewed through E_GSSG/2GSH_, have been closely related to loss of seed viability (Kranner *et al*., 2006; Kranner *et al*., 2010; Roach *et al*., 2010; Birtić *et al*., 2011; Chen *et al*., 2013; Morscher *et al*., 2015; Nagel *et al*., 2015; Roach *et al*., 2018a; Nagel *et al*., 2019;; Schausberger *et al*., 2019). However, in these studies seeds were likely aged at WCs above their Tg (i.e. with fluid cytoplasm). In *P. densiflora*, seed ageing was accompanied by shifts of E_GSSG/2GSH_ towards more oxidising cellular conditions, due to GSH depletion and GSSG accumulation (Figs. 2, 3A, B). Notably, hypoxia helped maintain more reducing cellular conditions compared to normoxia (Fig. 3B), indicating that O_2_ promoted ROS production also during seed ageing in the glassy state. Therefore, the redox conversion of GSH to GSSG and some non-enzymatic ROS scavenging by GSH were enabled within the highly viscous glassy cytoplasm.

Under normoxia, seeds aged at the lowest WC (0.004 g H_2_O g^-1^ DW, 9% RH) completely lost viability, despite their reduced cellular redox state (E_GSSG/2GSH_ = −195 mV; Fig. 3B). This value is more negative than the −180 to −160 mV range associated with a 50% loss of viability measured at higher seed WCs at 60% RH and 50 °C (Kranner *et al*., 2006). Reduced cellular redox states have also been found in unviable oil-rich seeds of *Vernonia galamensis* after ageing by CD in the glassy state, which contrasted to the more oxidised cellular redox states of seeds from the same species aged with fluid cytoplasm (Seal *et al*., 2010a; Seal *et al*., 2010b). The authors concluded that in this species E_GSSG/2GSH_ was less closely associated with viability after ageing by CD near or within the glassy state, agreeing with the results shown in the present study. Under dry/cold conditions of seed banks, seeds are typically in a glassy state, and the E_GSSG/2GSH_ values of barley seeds closely correlate to their viability after 15 years of seed bank-ageing (Nagel *et al*., 2015; Roach *et al*., 2018a). Similarly, *P. densiflora* seed bank seeds stored at low temperatures had only lost 13% of their viability, but their GSH concentrations were comparable to those detected in seeds aged by CD to complete viability loss under normoxia at 9% RH (Fig. 3A). Therefore, in glassy-state seeds temperature seems to influence O_2_-dependent deteriorative processes, which have down-stream consequences on GSH consumption. Indeed, during viability loss in the glassy state, seed bank seeds aged at low temperatures consumed more GSH than faster ageing seeds exposed to the higher temperature used for CD (Fig. 3A). However, it is important to consider that even if limited GSH consumption occurred while seeds were still desiccated, upon imbibition GSH concentrations may decrease following the GSTs-catalysed reactions with the abundantly produced RES (Fig. 6).

In summary, during seed ageing GSH consumption and redox conversion to GSSG were enhanced when the cytoplasm was fluid rather than glassy. However, these processes were not entirely restricted by the glassy state.

### A role for lipid peroxidation in the loss of viability of seeds with a glassy cytoplasm

Structural damage to cell membranes compromise solute compartmentalisation, leading to uncontrolled solute leakage and affecting cell functions (Powell and Matthews, 1981; Matthews and Powell, 2006). Normoxia in the glassy state resulted in cellular acidification (Fig. 3C), influencing the E_GSSG/2GSH_ values (Schafer and Buettner, 2001). In bread wheat, seed deterioration in the glassy state was accompanied by increases in the proton concentrations of seed extracts, explained as an effect of oxidative damage to the cell membranes (Nagel *et al*., 2019). Interestingly, *P. densiflora* seeds aged in the glassy state under normoxia leaked more electrolytes than seeds aged at the same RH under hypoxia (Fig. 3C), thus pointing to O_2_-mediated structural damage of cell membranes, likely implicated in the accelerated loss of viability.

Lipid peroxidation has been related to deterioration, particularly in oily seeds (Harman and Mattick, 1976; Pearce and Abdelsamad, 1980; Stewart and Bewley, 1980; McDonald, 1999; Tammela *et al*., 2005; Walters *et al*., 2005b; Oenel *et al*., 2017). However, also in starchy seeds of barley and wheat, oxidation and hydrolysis of TAGs and other lipids during ageing in the glassy state have been correlated with viability loss (Riewe *et al*., 2017; Wiebach *et al*., 2020). Furthermore, a decrease in the energy of lipid melting transitions, indicative of structural changes to the lipid phase, has been documented in aged seeds (Vertucci, 1992; Porteous *et al*., 2019). This phenomenon was also evident in CD-aged *P. densiflora* seeds under normoxia, but only after ageing in the glassy state (Fig. 4A) and can be explained by the depletion of unsaturated FAs, especially PUFAs (Fig. 4), which are more prone to peroxidation than unsaturated and monounsaturated FAs (Priestley and Leopold, 1983; McDonald, 1999; Smirnoff, 2010). The lipid melting peak L2 revealed by the DSC heating scans appeared at melting temperatures typical of the β’ crystals of linoleic (−25 °C) and linolenic (−35 °C) acids (Small, 1986; Knothe and Dunn, 2009), which were among the most abundant PUFAs of *P. densiflora* seeds (Fig. 1A, Supplementary Fig. S1) and have been found in other seeds and fern spores (Walters *et al*., 2005b; Ballesteros and Walters, 2007a;). However, the ΔH of lipid melt of peak L2 did not change after CD (Fig. 4A, Supplementary Fig. S1). In contrast, another lipid melting peak (L1) appeared at about −90 °C and sharply flattened in the DSC scans of seeds aged at 9% RH under normoxia (Supplementary Fig. S1). Depending on the cooling conditions, FAs can crystallise into different polymorphic types with the same chemical composition, but increasing order, density, and stability and decreasing energy and volume. These polymorphisms are generally denoted by the letters α, β’, and β, being α the first and least stable arrangement assumed by crystallising lipids (Metin and Hartel, 2005). The lipid melting peak L1 does not correspond to the melting temperature of β’ crystals of any tabulated TAG (Small, 1986; Knothe and Dunn, 2009), but likely resulted from the melting transition of α crystals of linoleic and linolenic acids, as observed in other seeds (Vertucci, 1992; Walters *et al*., 2005b). Therefore, it seems that peroxidation in the glassy state was mostly directed towards α crystals of linoleic and linolenic acids, contributing to peak L1 smoothing.

In the lipid domain of the cytoplasm, tocochromanols are the most abundant antioxidants essential to protect cells from lipid peroxidation and critical for seed quality (Menè-Saffranè *et al*., 2010). Recently, seed longevity has been associated with a high proportion of γ-tocopherol in the total vitamin E pool of several rice cultivars (Lee *et al*., 2020). Furthermore, seeds of tocochromanol-deficient mutants accumulate oxidised lipids and lipid-peroxide-derived RES, which lead to faster ageing (Sattler *et al*., 2004; Sattler *et al*., 2006; Menè-Saffranè *et al*., 2010). The presence of O_2_ during ageing of *P. densiflora* seeds in the glassy state resulted in a consumption of α- and γ-tocopherols (Fig. 5). This biochemical change in the lipid domain ties to increased electrolyte leakage during seed imbibition, changes in FA profiles, and drops in the ΔH of lipid melt (Figs 1C, 4), suggesting that O_2_ in the storage environment led to lipid peroxidation in seeds aged in the glassy state.

To ascertain the occurrence of lipid peroxidation, we measured peroxidation-associated products, including aldehydes and RES (Pamplona, 2011; Mano *et al*., 2019). The release of volatile aldehydes (e.g. hexanal) is a precocious symptom of lipid peroxidation during seed ageing in the glassy (WC < 0.05 g H_2_O g^-1^ DW) (Tammela *et al*., 2003; Mira *et al*., 2010), but not in the fluid state (Mira *et al*., 2016). Hexanal was a dominant aldehyde produced by *P. densiflora* seeds aged in the glassy state under normoxia (Fig. 6; Supplementary Fig. S4). Among the more reactive RES, 4-hydroxynonenal increased the most in response to CD of seeds with a glassy cytoplasm (Supplementary Fig. S5). Both these carbonyls are derived from ω-6 PUFAs, such as linoleic acid, whose contents significantly decreased in such seeds (Fig. 4B). Furthermore, PUFA-derived aldehydes can non-enzymatically convert to short-chain dicarboxylic acids (Passi *et al*., 1993). Indeed, azelaic acid, considered as a marker of lipid peroxidation in plants (Zoeller *et al*., 2012), increased 500-fold in seeds aged under normoxia at 9% RH compared to the control (Fig. 6). The C6 aldehydes (e.g. hexanal) can also be produced via lipid metabolism, involving lipoxygenase and hydroperoxide lyase during germination, but apparently not before imbibition (Weichert *et al*., 2002), supporting a non-enzymatic route of peroxidation-associated products’ formation during seed ageing in the glassy state. Therefore, the remarkably high concentrations of RES, aldehydes, and dicarboxylic acids detected under normoxia, confirmed that lipid peroxidation during ageing was strongly enhanced by O_2_ in glassy-state *P. densiflora* seeds (Fig. 6, Supplementary Fig. S4).

In summary, O_2_-mediated damage in the glassy state was characterised by deterioration of the seed lipid domain, the most mobile cytoplasmic domain in the glassy state. Loss of unsaturated FAs, enhanced production of RES and carbonyls, and consumption of tocopherols are all “hall-marks” of the O_2_-mediated autocatalytic cascade of lipid peroxidation (Fig. 7).

**Fig. 7.**
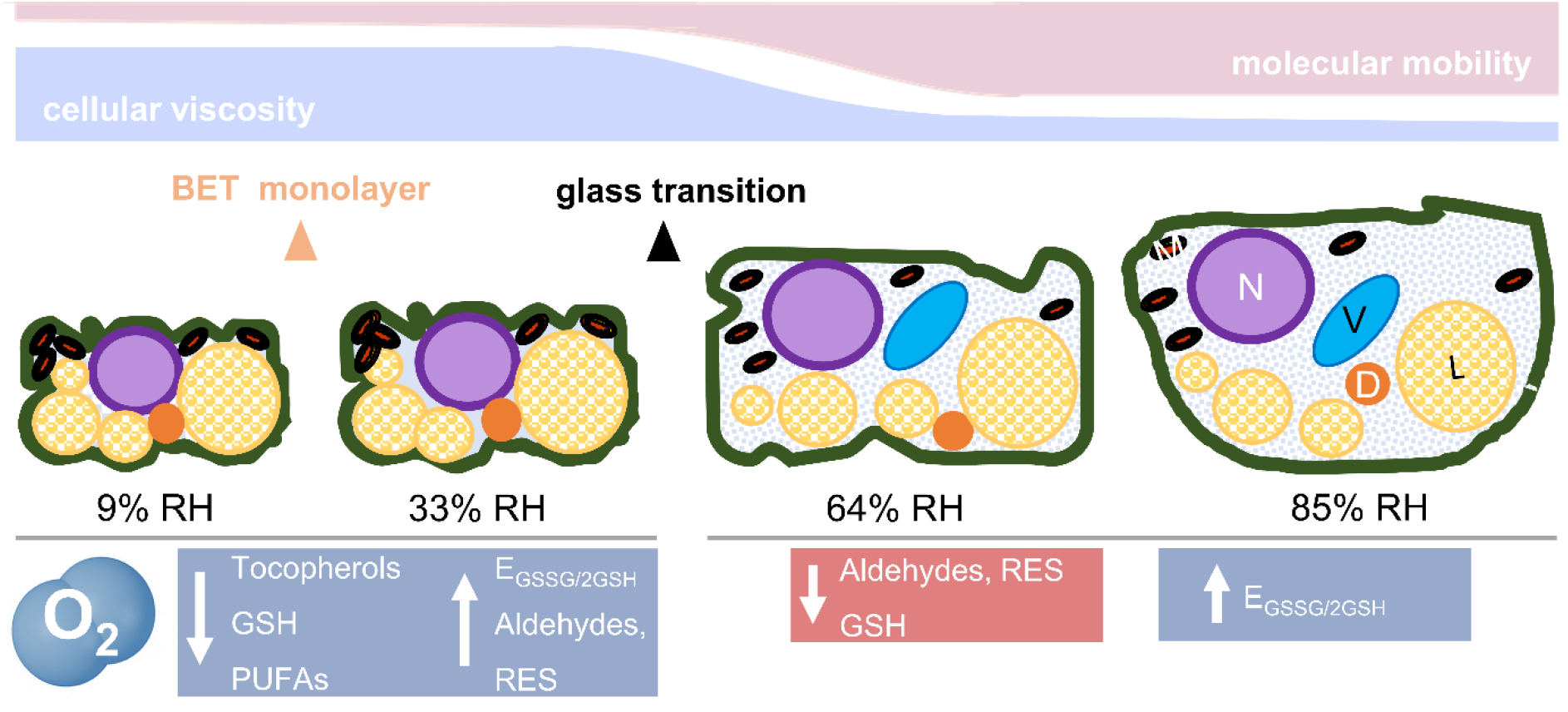
Schematic overview of the physicochemical changes of *Pinus densiflora* seed cells occurring in response to relative humidity (RH) and to O_2_ concentrations during controlled deterioration at 45 °C. During drying, the cytoplasm shrinks, reducing the area occupied by the cytosol and forcing in close proximity diverse organelles [nucleus (N), vacuole (V), mitochondria (M), and dry matter (D), including protein storage bodies and starch granules, the endomembrane system, and liquid lipid bodies (L)]. Cell walls and membranes are folded. Below a certain moisture content (< 0.05 g H_2_O g^-1^ DW or 42% RH, as from water sorption isotherms), the seed cytoplasm and organelles solidify (see Fig. 1B), forming an amorphous glass. Therefore, most of the seed cytoplasm at 9 and 33% RH (left side) was glassy. In addition, at 9% RH, the first monolayer of water molecules adsorbed to the surface of the macromolecules (i.e the Brunauer-Emmet-Teller [BET] monolayer) was partially removed, exposing some of their areas previously covered by water. In the glassy state, cellular viscosity is high, and molecular mobility is restricted to vibration, bending, and rotation of the side groups of macromolecules. In this physical state, O_2_ promoted lipid peroxidation, depletion of tocopherols, and accumulation of aldehydes and reactive electrophile species (RES). Between 42 and 50% RH (corresponding to 0.05 to 0.06 g H_2_O g^-1^ DW, see Fig. 1B), the cytoplasm changed from a glassy to a fluid state, which remains very viscous and is also known as “rubbery” state. This was the scenario for seeds aged at 64 and 85% RH (right side). In the fluid state, organelles tend to disperse due to the enlarged volume of the cytosol, and molecular mobility rises in comparison to the glass (i.e. the main chains of macromolecules are enabled to move). The biochemical changes indicated in blue were enhanced by O_2_, and those in red occurred independently of O_2_ availability. GSH, glutathione; PUFAs, (poly)unsaturated fatty acids; E_GSSG/2GSH_ half-cell reduction potential of the glutathione/glutathione disulphide redox couple. This figure is partly adapted from Ballesteros et al., 2020.

### Antioxidant metabolism resumes in rapidly-ageing seeds with fluid cytoplasm

In contrast to *P. densiflora* seeds aged by CD in the glassy state, the longevity of those seeds aged by CD with fluid cytoplasm (i.e. at 64 and 85% RH) was not extended by hypoxia, and no significant signs of lipid peroxidation were detected (Figs. 2, 4B). This agrees with the release of volatiles by seeds aged with fluid cytoplasm, as reported in previous studies, which also pointed to oxygen-independent glycolytic and fermentations reactions (Mira *et al*., 2010; Colville *et al*., 2012). Previous analyses on sunflower, barley, and broccoli suggest that elevated O_2_ concentrations are not detrimental to the longevity of seeds aged by CD with fluid cytoplasm (Morscher *et al*., 2015; Roach *et al*., 2018a; Schausberger *et al*., 2019). However, in these studies the modulation of O_2_ during CD affects the concentrations of LMW antioxidants. For instance, various tocochromanols increase in response to CD, but differently depending upon O_2_ availability (Roach *et al*., 2018a). This result aligns to the finding that enzyme activity, which reinforces antioxidant defences, is possible in the “rubbery” (fluid) state, but not in the glassy state (Fernández-Marín *et al*., 2013; Candotto Carniel *et al*., 2021). Whereas the majority of steps in tocopherol synthesis occurs within the lipid phase of the cytoplasm, precursors (e.g. tyrosine), intermediates, and substrates for the pathways (e.g. ATP) are located in the aqueous domain and necessitate sufficient molecular mobility to be accessible to enzymes (Menè-Saffranè and DellaPenna, 2010; Muñoz and Munné-Bosch, 2019). Conversely, *de novo* GSH biosynthesis takes place entirely in the aqueous domain and requires two ATP-dependent reactions, the first of which is the rate limiting step and generates γ-Glu-Cys at the expense of ATP (Noctor *et al*., 2012). Therefore, increases in γ-Glu-Cys concentrations in *P. densiflora* seeds aged at 64% and 85% RH could indicate GSH anabolism (Supplementary Fig. S3). Alternatively, other enzymes (e.g. carboxypetidases) could account for the release of γ-Glu-Cys during GSH catabolism (Noctor *et al*., 2012). The ligase that catalyses γ-Glu-Cys formation (EC 6.3.2.2) is regulated by GSH and Cys concentrations via non-allosteric feedback competitive inhibition with glutamate (Yang *et al*., 2019). Consequently, the depletion of GSH could have stimulated GSH *de novo* synthesis, as part of protective antioxidant mechanisms. In fact, redox homeostasis ensured by GSH availability also prevents RES from being highly toxic molecules (Farmer and Mueller, 2013).

One route that could lead to GSH depletion, rather than GSSG accumulation, relies on GST-mediated conjugation to RES. Some RES, such as acrolein, have profound impact on GSH concentrations (Mano, 2012; Roach *et al*., 2018b), and in the present study the concentrations of RES decreased in seeds aged with fluid cytoplasm (Fig. 6), conditions in which GSH concentrations dropped most (Fig. 3A).

In summary, our data suggest that enzymatic activity can resume in seeds with a fluid cytoplasm, whereby damage to lipids appeared to be very marginal, differently from glassy-state seeds. Besides glycolysis and fermentation reactions, reported in previous studies, here we show that seeds in the fluid state also resumed a certain level of antioxidant metabolism (Fig. 7), which may include γ-Glu-Cys synthesis and GST activity, potentially to counteract rapid ageing rates.

### Ageing by CD in the glassy state best simulates biochemical changes during cold seed storage

After storage at WC of ~0.06 g H_2_O g^-1^ DW for 20 years at temperatures between +4 and −20 °C, *P. densiflora* seeds contained less GSH and γ-tocopherol and showed signs of lipid peroxidation, such as elevated concentrations of hexanal and 4-hydroxynonenal, less unsaturated FAs, and more pronounced electrolyte leakage in comparison with non-aged control seeds (Figs. 2–6, Supplementary Fig. S4). We cannot exclude that environmental conditions of the different harvest years contributed to some biochemical differences between seed bank seeds (1999) and seeds used for CD (2015). Nonetheless, this comparison revealed biochemical changes under low storage temperatures consistent with changes induced by CD when conducted in the glassy but not in the fluid state.

### Implications for seed longevity prediction and germplasm storage

The differential mechanisms of seed ageing found between seeds in the glassy and the fluid state contribute to the debate about the use of CD or accelerated ageing methods in seed science research, but also highlight the potential benefits of hypoxic dry seed storage in seed banks to extend longevity. Equations that include temperature, RH, species-specific ageing constants, and initial seed viability have been derived for estimating seed ageing rates, but as yet they do not include the influence of O_2_ concentration (Ellis and Roberts, 1980; Pritchard and Dickie, 2003; Ellis and Hong, 2007). This would be highly relevant considering that in seed banks, germplasm is typically stored in the glassy state and, in some cases, is ageing faster than expected (Li and Pritchard, 2009). Therefore, our results have three main implications for seed banking management. Firstly, for studying seed longevity, CD better reflects ageing mechanisms under the dry/cold conditions of a seed bank when seeds are treated in the glassy rather than fluid state. Secondly, as O_2_ promoted seed ageing reactions during CD below the Tg, limiting seed exposure to O_2_ during long-term cold storage, as also recommended by other authors (Groot *et al*., 2015; Nagel *et al*., 2016; Buijs *et al*., 2020) would most likely prolong germplasm longevity. Finally, equations to predict seed longevity under cold, dry, and hypoxic conditions require new viability constants to be calculated.

## Abbreviations

AsA: ascorbic acid;
BET: Brunauer-Emmet-Teller;
CD: controlled deterioration;
Cys: cysteine;
Cys-Gly: cysteinyl-glycine;
ΔH: enthalpy;
DMA: dynamic mechanical analyses;
DNPH: 2,4-dinitrophenylhydrazine;
DSC: differential scanning calorimetry;
DTT: dithiothreitol;
DW: dry weight;
EC: electrical conductivity;
E_GSSG/2GSH_: half-cell reduction potential of the glutathione/glutathione disulphide redox couple;
E_hc_: half-cell reduction potential;
*E*^0^_pH_: standard half-cell reduction potential at a defined pH;
FA: fatty acid;
FAME: fatty acid methyl ester;
FW: fresh weight;
γ-Glu-Cys: γ-glutamyl-cysteine;
GC-MS: gas chromatography coupled to mass spectrometry;
GSH: glutathione;
GSSG: glutathione disulphide;
GST: glutathione-S-transferase;
HPLC: high-performance liquid chromatography;
LMW: low-molecular-weight;
P50: time to decrease seed viability by 50%;
PUFA: polyunsaturated fatty acid;
RES: reactive electrophile species;
RH: relative humidity;
ROS: reactive oxygen species;
RT: room temperature;
T_25_: time to reach 25% germination;
TAG: triacylglycerols;
TD-NMR: time-domain nuclear magnetic resonance;
Tg: glass transition temperature;
uHPLC-MS/MS: ultra-high performance liquid chromatography tandem mass spectrometry;
UPW: ultrapure water;
WC: water content.

## Supplementary data

Supplementary data are available at *JBX online*.

*Table S1*. Conditions used for controlled deterioration (CD) of *Pinus densiflora* seeds under normoxia and hypoxia.

*Fig. S1*. Effects of relative humidity (RH) on the influence of O_2_ during controlled deterioration (CD) at 45 °C on the melting properties of seed storage lipids, measured with differential scanning calorimetry (DSC).

*Fig. S2*. Seed longevity after controlled deterioration (CD) at 45 °C under normoxia (i.e. 19.6% O_2_) and RH ranging from 30% to 100%, corresponding to indicated seed water contents (WCs) determined after lyophilisation.

*Fig. S3*. Effects of relative humidity (RH) on the influence of O_2_ during controlled deterioration (CD) at 45 °C on seed germination.

*Fig. S4*. Effects of relative humidity (RH) on the influence of O_2_ during controlled deterioration (CD) at 45 °C on total concentrations of low-molecular-weight thiol/disulphide redox couples.

*Fig. S5*. Effects of relative humidity (RH) on the influence of O_2_ during controlled deterioration (CD) at 45 °C on absolute concentrations of reactive electrophile species and aldehydes.

## Acknowledgments

The authors thank Birgit Knoll and Bettina Lehr (University of Innsbruck) for excellent technical assistance with oxygen and electrical conductivity measurements and HPLC analyses.

Funding by the Ministry of the Republic of South Korea to CSN are gratefully acknowledged (NRF-2017R1D1A1B03034615). The BritInn Fellowship Program (Academic Network Britain Innsbruck, University of Innsbruck) supported DG with a research travel grant to the Royal Botanic Gardens, Kew. The Royal Botanic Gardens, Kew, receive grant-in-aid from DEFRA.

## Author contributions

Conceptualisation: TR, DG, DB; formal analysis: DG, DB, WS, EA, TR; funding acquisition: TR, CSN, DG; investigation: DG, TR, DB, CES, CSN; methodology: DG, TR, DB, WS, EA, CES; project administration: TR; resources: CSN, CES, IK; supervision: TR, DB, CES; validation: DG, TR, DB, WS, EA, CES; visualisation: DG, TR, DB; writing - original draft: DG, TR, DB; writing - review & editing: TR, DG, DB, WS, EA, CES, IK. All authors read and approved the final version of the manuscript.

## Data availability statement

The data supporting the findings of this study are available from the corresponding author, TR, upon request.

